# A Morphogenetic Wave that Generates Mesenchymal-to-Epithelial Transition in the Lateral Plate Mesoderm

**DOI:** 10.1101/2022.03.01.482468

**Authors:** Manar Abboud Asleh, Mira Zaher, Julian Jadon, Lihi Shaulov, Ronit Yelin, Thomas M. Schultheiss

## Abstract

Most mesodermal cells undergo multiple cycles of transition between an epithelial and mesenchymal state during embryonic development. While many studies have addressed the process of epithelial-to-mesenchymal transition (EMT), comparatively less is known regarding the complementary process, mesenchymal-to-epithelial transition (MET), which is essential for organogenesis and has also been proposed to be important for cancer metastasis. The current study investigated MET using the lateral plate mesoderm (LPM) of the chick embryo as a model system. We find that MET in the LPM proceeds as a wave, which divides the LPM into distinct mesenchymal, transition, and epithelial zones. In the multilayered mesenchymal zone, many apical epithelial markers, including N-Cadherin (N-Cad), Par-3 and Zo-1, but not atypical protein kinase C (aPKC), are detected as dispersed, partially co-localizing aggregates associated with cell-cell contacts. The transition zone is characterized by the appearance of aPKC and the formation of rosette-like structures characterized by wedge-shaped cells that are apical-basal polarized, with strong co-localization of apical polarity markers, but not yet arranged into distinct epithelial sheets. The transition zone is also enriched in mitotic cells. Subsequently, the rosettes resolve into two well-defined epithelial sheets that constitute the coelomic epithelium, the lining of the internal body cavity.

Prior to any overt signs of apical-basal polarity, fibronectin (FN) begins to accumulate at the future basal side of the incipient epithelium. Interference with Extracellular Matrix (ECM)-integrin signaling through disruption of focal adhesion kinase (FAK) or Talin function hindered the normal progression of the epithelialization process. Cells with disrupted FAK or Talin function retained mesenchymal-like characteristics with respect to cellular morphology and apical-basal marker distribution.

We propose a two-stage process for MET in the LPM. Initially, in the polarization phase, ECM-integrin-dependent signaling imparts apical-basal polarity, culminating in the activation of aPKC, to drive cell intercalation and rosette formation. Subsequently in the resolution phase, polarized rosette cells, perhaps facilitated by the weakening of cell-cell interactions that occurs during mitosis, expand their apical surface, and spread out to form new connections laterally to their fully epithelial neighbors. This sequence of events is propagated as a wave through the LPM, thus generating an integrated coelomic epithelium.

## Introduction

Mesenchymal-to-Epithelial Transition (MET) is a fundamental biological process in which individual mesenchymal cells become organized into epithelial sheets^1,2^. In the embryo, MET is critical to the formation of somites, kidney tubules and ducts, the coelomic lining of the body cavity, and other mesodermally-derived tissues^3–7^. MET has also been reported to be important during cancer metastasis, where it has been implicated in the ability of metastatic cells to establish tumors in distant sites^8–10^. In addition, the process of wound healing in the skin and other organs involves first a stage in which epithelial cells lose their epithelial properties and spread to cover the wound, followed by a stage of re-epithelialization to restore the epithelial barrier of the wounded tissue^11,12^. Defects in MET are implicated in a number of pathologies, including kidney malformation and lumenopathies^2,13^.

MET is often considered together with its converse and better-studied sister process Epithelial-to-Mesenchymal Transition (EMT), in which cells acquire migratory properties and leave an epithelial sheet^14^. EMT is essential to fundamental embryological processes including gastrulation, neural crest migration, and limb bud formation^15–17^, as well as the spread of cancer metastases^18^. Recent analysis has emphasized that “Epithelial” and “Mesenchymal” are not absolute states, but rather poles of a spectrum (the Epithelial-to-Mesenchymal Spectrum, or EMS)^19,20^. For example, cells undergoing Collective Cell Migration (CCM), such as neural crest cells, kidney duct, or lateral line organ, exhibit both epithelial (cohesion) and mesenchymal (migratory) properties that are essential for their biological functions^21–23^.

Epithelial cells are polarized, with distinct apical, lateral and basal domains and asymmetric distribution of proteins and cellular organelles within the cell^1,24–26^. In many systems, the apical complex includes Crumbs, Par-6, and atypical protein kinase C (aPKC), whereas Par-3 (bazooka) localizes to the apex of the lateral domain, and the basolateral complex is characterized by localization of Disc Large (Dlg), Scribble, Lethal giant larvae (Lgl) and Par-1. This specific localization of the different complexes is maintained by mutual repressive and collaborative interactions, and by the presence of adherens junctions, which strongly interconnect the epithelial cells. On the basal side, epithelial cells express integrin and dystroglycan receptors that interact with the extracellular basement membrane and extracellular matrix (ECM).

Many studies have identified regulators of EMT including signaling, transcriptional, and biomechanical factors^14,15,27^. In contrast, less is understood regarding the molecular regulation of the MET process. MET requires that mesenchymal cells aggregate, acquire apical-basal polarity, and form mature cell-cell junctions that join cells to their neighbors. Studies in the early mouse embryo have pointed to the ECM, and in particular laminin (LN), as providing external cues to orient the polarity of egg cylinder cells undergoing epithelialization^28^. ECM is also required for epithelial tube formation in MDCK cells^29^, a frequently studied in-vitro model for epithelialization. In vertebrate kidney formation, Wnt signaling is required for the transition of metanephric mesenchyme cells to renal vesicles, the epithelial precursors of kidney tubules^4,6,30,31^. Embryonic somites are also formed through MET; the segmentation clock that regulates partitioning of the paraxial mesoderm into somites has been extensively studied^32,33^, and some regulators of MET in the paraxial mesoderm have been identified^34^. However, despite this progress in several experimental systems, a mechanistic understanding of the stages of MET as it occurs in vivo is still far from complete.

The current study makes use of the chick embryo lateral plate mesoderm (LPM) as an in vivo model system for studying MET. The LPM is formed by newly-gastrulated mesenchymal cells that derive from the posterior half of the primitive streak^35,36^ and migrate to lateral regions of the embryo. These cells subsequently undergo MET to generate the coelomic epithelium (Co-Ep) (Fig. 1). The Co-Ep consists of two pseudostratified epithelial layers, the splanchnopleuric (Spl) on the endodermal side, and the somatopleuric (Sop), on the ectodermal side^37^. The apical surfaces of the Spl and Sop face each other and enclose a fluid-filled space called the coelom, or body cavity. Subsequently, portions of the Co-Ep undergo EMT to generate the smooth muscle and mesenteries of the gut, the mesenchyme of the gonads, the limb bud, and other structures, while the remaining Co-Ep cells constitute the permanent epithelial lining of the body cavity, including the peritoneum, pleura, and pericardium^16,37,38^. The current study describes MET in the LPM in morphological and molecular detail, identifying a novel morphogenetic wave propelled by ECM-integrin signaling that sweeps across the embryo and organizes the mesenchymal cells into an epithelium through a rosette-like-transition structure.

**Figure 1.**
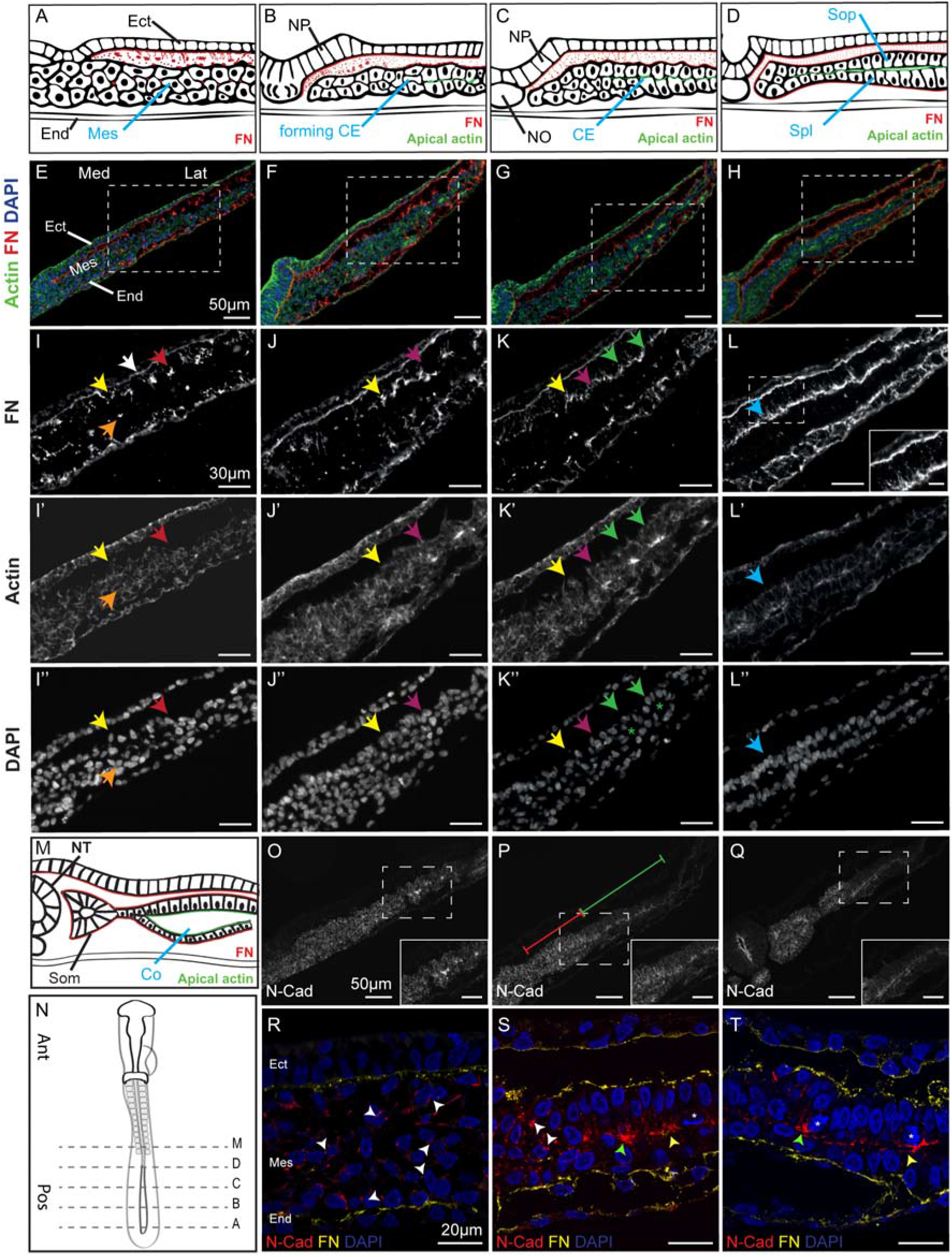
Wavelike progression of MET in the Lateral Plate Mesoderm. (A-L) The four columns (A,E,I; B,F,J; C,G,K; and D,H,L) depict four successive stages in MET. In each panel, the right half of the embryo is shown, and medial is on the left. The boxed areas in EH are magnified in I-L. In I-K, yellow arrows indicate the most medial extent of FN accumulation on the basal side of the Sop, magenta arrows indicate the most medial apical actin enrichment, and green arrows indicate rosettes. Note that the basal FN always extends more medial than the apical actin localization and the rosettes. White arrow indicates ectodermal FN, red arrows indicate punctate FN, and orange arrows indicate FN in the interior of the mesoderm. The boxed area in L is shown at higher magnification in the inset and it demonstrates the fibrillar FN extending between the ectoderm and mesoderm (the scale bar is 10μm). Blue arrows point to area where the coelomic cavity has begun to open. (M) Subsequent stage in MET showing opening of the coelom. (N) Approximate axial levels of indicated cross sections in a Stage 10 embryo. (O-Q) Progression of N-Cadherin (N-Cad) distribution during MET as seen by widefield microscopy. Note the transition of N-Cad staining from dispersed (red segment in O) to apically localized (green segment) N-Cad. The boxed areas are shown at higher magnification in the insets (the scale bar is 30μm). (R-T) Higher magnification confocal views showing N-Cad and FN distribution at three stages of MET. White arrowheads indicate dispersed N-Cad staining. Green arrowheads indicate rosettes. Yellow arrowhead indicates apical concentration of N-Cad upon epithelialization. Asterisk indicates mitotic cells. FN, fibronectin; N-Cad, N-Cadherin; CE, coelomic epithelium; Ect, ectoderm; End, endoderm; Mes, mesoderm; NO, notochord; NP, neural plate; NT, neural tube; Co, coelom; Som, somite; Sop, somatopleuric mesoderm; Spl, splanchnopleuric mesoderm.

## Results

### MET proceeds as a morphogenetic wave within the lateral plate mesoderm

In initial studies, we examined the process of MET in the lateral plate mesoderm (LPM) of HH Stage 9 to 11 embryos using a set of molecular markers to visualize cell boundaries and cellular organization and to characterize apical and basal subcellular domains (Fig. 1). At HH Stage 9, LPM cells in the middle to posterior region of the embryo exhibit a typical mesenchymal morphology (Fig. 1A,E,I-I”). Actin is uniformly distributed throughout the LPM, and nuclei exhibit a dispersed distribution. At this stage there are already hints of incipient polarity, with Fibronectin (FN) aggregates enriched on the future basal side of the LPM (Fig. 1I red arrow), although FN aggregates are also detected in the interior of the LPM layer (Fig. 1I orange arrow). These basal FN aggregates appear rudimentary compared to the mature, continuous FN layers of the ectodermal and endodermal basement membranes (Fig. 1I white arrow).

Subsequently, signs of polarization and epithelialization progressively appear within the LPM. As described in detail below, these changes appear first in lateral, anterior regions of the LPM and then spread in a wave into medial and posterior regions (i.e. in a diagonal fashion). Because of this wave-like progression of MET, a cross-section of an embryo that is in the midst of these changes contains concomitantly regions that exhibit all stages of the epithelialization process, with more advanced stages laterally and less advanced stages medially, which greatly aides in characterizing the timing of events in the MET process. In the current description and in the figures, we emphasize events occurring in the Sop layer of the LPM; however similar events occur also in the Spl layer.

At the leading edge of the MET wave, FN aggregates elongate and integrate into a more continuous layer on the future basal side of the Co-Ep (Fig. 1J, yellow arrow) (Fig. 1K-L). Lagging slightly more laterally, discrete foci enriched in actin are detected in the middle (in the dorsal-ventral dimension) of the LPM (Fig. 1J’-K’, green arrows), which also spread from lateral to medial. These actin-containing foci are at the center of rosette-like cellular arrangements (Fig. 1K’,K”, green arrows and asterisks; see also Figs. 2,3). Subsequently, the rosette-like structures resolve into two polarized epithelial sheets, the somatic (Sop) and splanchnic (Spl) layers of the Co-Ep (Fig. 1D,H,L-L”), which are outlined by strong basal FN accumulation (Fig. 1L) and lined by prominent apical actin concentration (Fig. 1L’). At these stages, intra-mesodermal FN can no longer be detected. Thereafter, slits appear between the epithelial sheets (Fig. 1L’, blue arrow), which later coalesce to form the coelomic cavity (Fig. 1M). At this stage, the negatively charged sialomucin podocalyxin^28,39^ begins to be detected, lining the coelomic slits (Sup. Fig. S1A-F).

**Figure 2.**
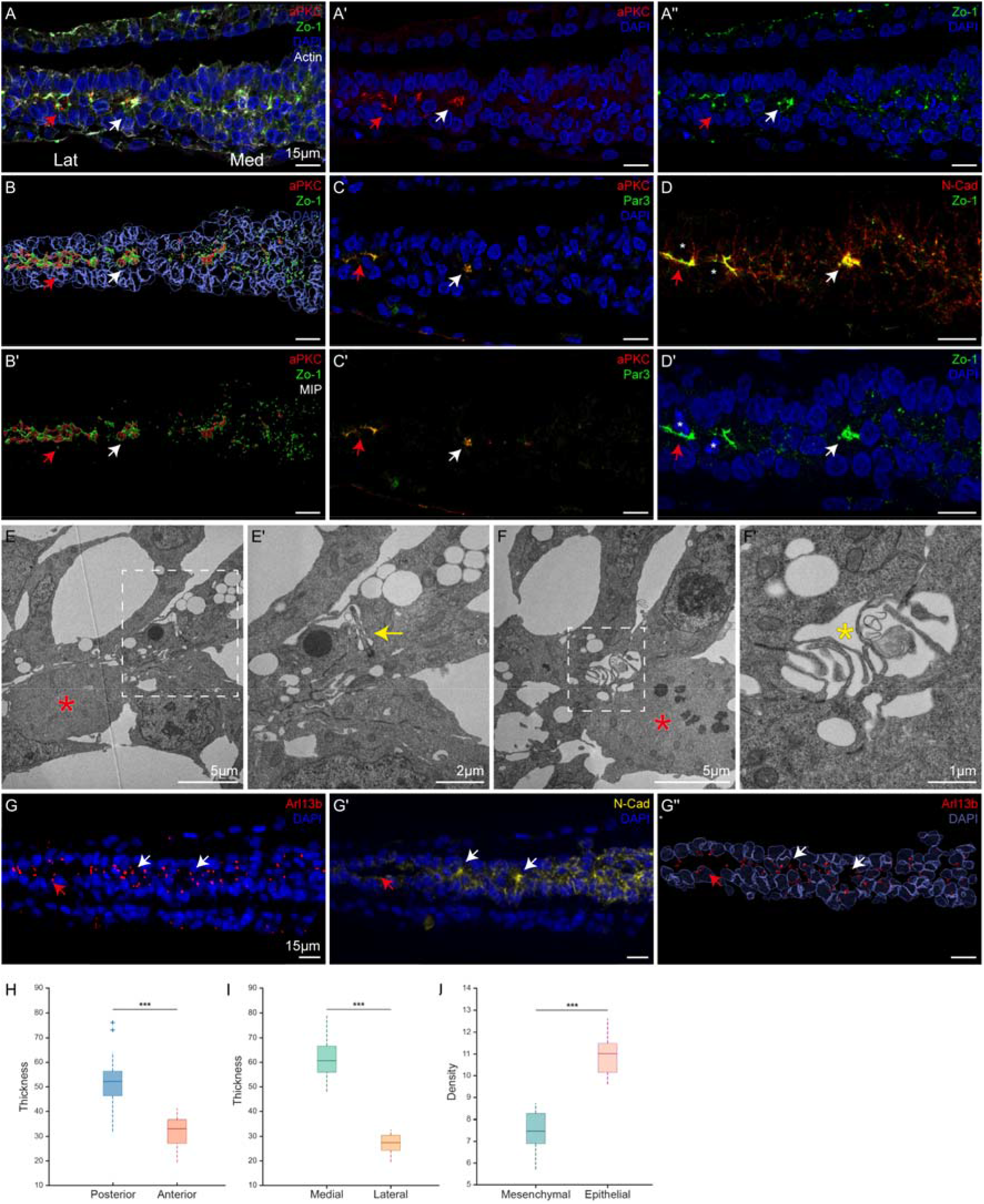
Molecular and Morphological Characterization of Rosette Formation during MET. Medial is on the right. (A-D,G-G”) Expression of apical markers during MET. White arrows indicate rosettes, and red arrows indicate epithelium. Zo-1 (A,A”,B,B’,D,D’) undergoes reorganization from a dispersed distribution in mesenchymal regions to central aggregation in rosette complexes and apical distribution in the nascent epithelium; aPKC (A,A’,B,B’C,C’) is not detected in the mesenchyme and appears apically in the rosettes and epithelium; Par-3 (C,C’) undergoes upregulation and reorganization in the transition from mesenchyme to epithelium; and N-Cad (D,G’) starts off as a network of dispersed dots around the cell membrane at the mesenchymal stage (medial side), and then gradually relocalizes to the apical side in the rosettes and epithelium. Asterisks indicate mitotic cells. B,B’ are a model constructed from 3D projections from a segmented confocal Z-series, with the nuclei rendered semi-transparent. Note the clear demarcation of mesenchymal, transition (rosette) and epithelial regions, and Zo-1 and N-Cad expression. (E,F) Electron Microscopy of rosette centers showing triangular/wedge-shaped cells with apical cilia (E’ yellow arrow) and extensive apical junctions. (F,F’) Shows the first ultrastructural stage of rosette-resolution with multiple apical membrane folds extending into the small central lumen (yellow asterisk). Red asterisks in E,F indicate mitotic cells. See Sup. Fig. S2 for lower power views. (G-G”) Cilia, as marked by Arl13b, undergo reorganization from a random to an apical distribution during MET. G” is a 3D-model constructed from a confocal Z-series, as in B. (H-I) Comparison of mesodermal thickness in anterior vs. posterior (H) and medial vs. lateral (I) regions. (J) Comparison of nuclear density in mesenchymal vs. epithelial regions. The mesodermal layer thins and increases in density as MET progresses.

**Figure 3.**
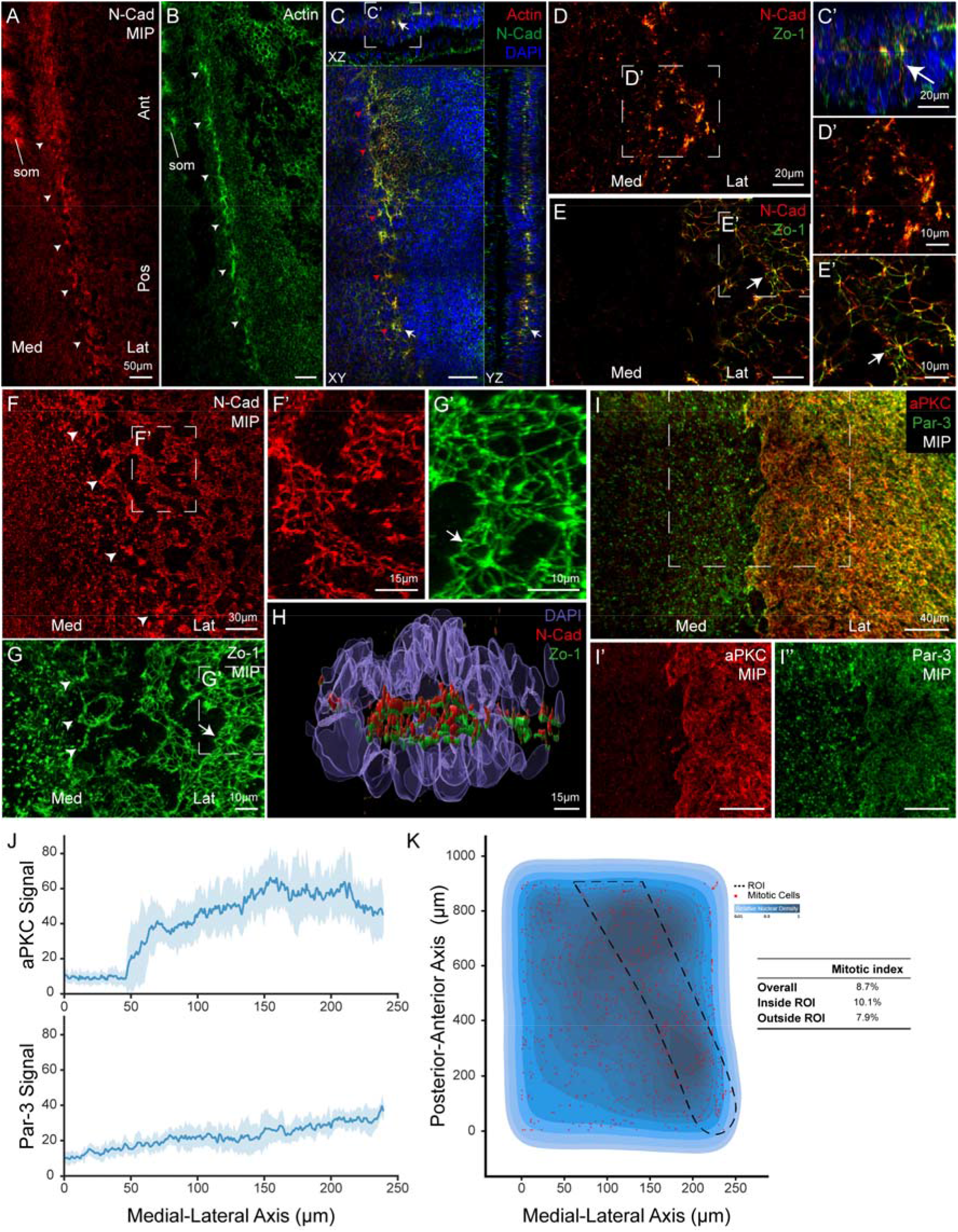
Characterization of the Line of Rosettes at the Forefront of Epithelialization. Medial is on the left. (A-B) Whole mount flat (XY) view of embryo stained for N-Cad (A in Multiple Intensity Projection [MIP]) and Actin (B, single plane) showing the line of rosettes (arrowheads). (C) Whole mount flat (XY) view and XZ and YZ projections of embryo undergoing MET. White arrow indicates rosette that can be seen in all planes (enlarged in C’). Note the line of rosettes (red arrowheads). (D-E) Whole mount flat XY view of Zo-1 and N-Cad stained-embryo undergoing MET. D and E are individual Z sections from the same Z-stack. D’ and E’ are magnifications of mesenchymal (medial) and epithelial (lateral) regions, respectively. These regions appear on separate Z-levels because of differences in morphology of the embryo between medial and lateral regions and because the embryo is slanted in the microscope on the XZ axis. Note the honeycomb-like staining in E’ due to viewing of the apical staining en face compared to the dotted dispersed pattern of the mesenchymal stage. F and G are MIPs of the same image stack as D-E, and F’ and G’ are magnified views of the indicated areas. Arrows in E,E’,G,G’ indicate “hemi-rosettes” as seen en face, from inside the coelomic space. Arrowheads in F, G indicate the line of rosettes. (H) 3D Model of a single rosette segmentation from this data set. (I) Whole mount flat (XY) view of embryo stained for aPKC and Par-3, viewed in MIP. Note the transition in Par-3 from dispersed to honeycomb-like pattern (I’’) and the abrupt emergence of aPKC upon initiation of epithelialization (I’). (J) Quantification of signal intensity for aPKC and Par-3. (K) Distribution of mitoses in the mesoderm. Heat map of proportional cell density is shown in blue, and mitoses are indicated as red dots. The black border indicates the region of interest (ROI) enclosing the line of rosettes. Note increased mitotic density in the vicinity of the line of rosettes.

The wave-like nature of MET progression is seen particularly clearly when analyzing expression of the cell-cell adhesion protein N-Cadherin (N-Cad; LPM cells express N-Cad and not E-Cad^40^). In the mesenchymal state (Fig. 1O, R, red bracket in S), N-Cad is dispersed throughout cellular membranes, predominately at cellular contacts (Fig. 1R,S white arrowheads). Gradually, N-Cad relocalizes and concentrates at apical foci of rosette-like structures (Fig. 1P green bracket, S green arrowhead), which subsequently resolve into epithelial sheets with strong apical “zipper-like” expression of N-Cad (Fig. 1Q,T).

### Molecular and Morphological Characterization of Rosette Formation and Resolution during MET

In order to study more closely the MET transition, sectioned embryos were examined with a broader set of epithelial polarity markers and also ultrastructurally (Fig. 2, Sup. Fig. S2). The tight junction component Zo-1 and the apical-basal polarity organizer Par-3 are expressed in mesenchymal regions in a dispersed pattern (Fig. 2A,A”,B,B’,D,D’, and C,C’ respectively) that largely colocalizes with N-Cad (Fig. 2D), although N-Cad is more broadly distributed. Similar to N-Cad, Zo-1 and Par-3 also undergo relocalization to apical foci in rosettes (Fig. 2A,B,D and C,C’, respectively, white arrows) and to the apical face of the newly-formed epithelium (Fig. 2B,D and C, C’, respectively, red arrows). Par-3 is expressed weakly but detectably in mesenchymal regions (Fig. 2C’ right side; see Fig. 3I,I” for stronger signal using Maximal Intensity Projection). In contrast, the polarity protein aPKC is almost undetectable in the mesenchymal zone, and appears suddenly and strongly in the apical regions of the rosettes and epithelium (Fig. 2A’, B,B’,C,C’, see also Fig. 3I,I’).

Examination of the transition region with Transmission Electron Microscopy (TEM) reveals that rosettes are comprised of triangular or wedge-shaped cells. These cells exhibit well-developed apical domains, including extensive junctional complexes and apically-oriented cilia (Fig. 2E,E’,G,G’,G’’; Sup. Fig. S2E-H). This is in contrast to the multilayered mesenchymal regions, where cells form focal contacts with each other (Sup. Fig. S2 A-D red arrows). Incipient ultrastructural rosette resolution is characterized by separation between the apical surfaces, forming a small central space containing abundant membrane protrusions (Fig. 2F,F’; Sup. Fig. S2I-K). Eventually, these membranous extensions are shed (Sup. Fig. S2M-P), suggesting active rearrangement of the apical membrane domain in the rosettes. Subsequently “hemi-rosette” morphologies, can be observed, which consist of intralayer apically-joined wedge-shaped cells (Sup. Fig. S2Q-T, Sop layer), separated by a small space from the opposing layer that no longer exhibits a rosette-like morphology.

In addition to the molecular and ultrastructural changes happening during MET, gross morphological changes are also noticeable. Consistent with the cellular observations showing cell aggregation and intercalation during MET progression (Fig. 1B,F,J-J”, 2A-D’,G-G”), measurements of LPM thickness and cell density revealed that the lateral and anterior (more epithelial) regions were significantly thinner and denser than the medial and posterior (mesenchymal) regions (Fig. 2H-J; see also Fig. 3K). This indicates that the multilayered mesenchyme undergoes extensive compaction to transform into a two-layered epithelium and raises the possibility that the LPM may undergo a type of convergent extension during MET, wherein the layer converges (thins) in the dorsal-ventral dimension as it extends in medial-lateral and anterior-posterior axes^40^.

In summary, three zones can be distinguished in the LPM that is undergoing MET: a mesenchymal zone characterized by a loose arrangement of multilayered mesenchymal cells and dispersed intracellular distribution of apical-basal markers; a transition zone characterized by the aggregation and intercalation of cells into rosette-like structures that exhibit apical-basal polarity, apical junctional complexes, and the de novo appearance of aPKC; and an epithelial zone in which the rosette structures resolve into two mature epithelial layers.

### A Line of Rosettes Guides the MET Transition

The results thus far were derived from observations on sectioned embryos. In order to obtain a more comprehensive view of the MET process in the entire LPM, we examined confocal Z-stacks of whole, cleared, immunostained embryos. This method allowed us to study and understand the morphology and role of the rosette-like structures that are at the forefront of the epithelialization process.

These studies revealed a diagonal line, intermittent posteriorly and more continuous anteriorly, of enriched foci of apical and junctional markers, including actin and N-Cad (Fig. 3A-C arrowheads), as well as Zo-1, aPKC, and Par-3 (data not shown). Actin cables connect the foci along this line (Fig. 3B,C). Virtual cross-sections reveal that the foci correspond to the rosettes described in Fig. 2 (Fig. 3C,C’), and that these rosettes are three-dimensional (Fig. 3H and supplemental videos S1,2). Hereafter, this line will be referred to as the *line of rosettes*. Medial to this line, the cells maintain a mesenchymal phenotype, while laterally the mesodermal layer is largely epithelial. In flat-mount (XY plane) view, this line separates the dotted, dispersed appearance of the various apical markers (N-Cad, Zo-1, and Par-3) medially from their honeycomb-arrangement laterally, which represent the apical side of the epithelial sheets (Fig. 3D-I, supplemental video S3). Within the hexagonal pattern of N-Cad and Zo-1, flat hemi-rosette structures can be seen (Fig. 3E,E’, white arrow), which correspond to the hemi-rosettes seen in the EM observations (Sup. Fig. S2Q-T), but here viewed en face (i.e. in XY view, as seen from inside the coelomic slits). In agreement with what was observed in sections, the apical marker aPKC undergoes the most dramatic transition, from largely undetectable in the mesenchymal zone to strong expression in the apical regions of cells in the transition and epithelial regions (Fig. 3I,I’).

Confirming the observations in Fig. 3I, global signal intensity quantification revealed a gradual medial-to-lateral increase in Par-3 expression and an abrupt and steep increase in aPKC expression at the line of rosettes (Fig. 3J). In contrast, N-Cad (see Fig. 6O, red line) and Zo-1 intensity (data not shown) were essentially uniform across the medial to lateral axis, indicating that their changes can be attributed to changes in protein localization as opposed to changes in expression levels. This suggests that Par-3 and especially aPKC, may play a role in reorganizing pre-existing apical and junctional complex in mesenchymal cells into a polarized, epithelial arrangement. aPKC mRNA was not significantly upregulated in lateral as opposed to medial zones (Sup. Fig. S3A-C), indicating that the increase in aPKC staining intensity is post-transcriptionally regulated.

Colocalization analysis revealed strong colocalization between Zo-1 and N-Cad throughout the MET process (supplemental videos S1,2,3). Most Zo-1 expression overlaps with N-Cad, especially medially (Manders coefficient (MC) 0.98 medially, 0.88 laterally), although N-Cad expression is much broader, particularly medially (MC 0.18 medially, 0.48 laterally). Par-3 shows significant colocalization with N-Cad medially (MC 0.69), which decreases laterally (MC 0.5). As with Zo-1, N-Cad is more broadly expressed than Par-3 (MC 0.34 and 0.48, respectively). As discussed above, aPKC is virtually undetectable in medial regions. When aPKC appears in the rosette and lateral regions, it is well-correlated with Par-3 (MC 0.67). Together these data indicate significant colocalization of Zo-1, Par-3, and N-Cad throughout the MET process, with the addition of co-localized aPKC at the onset of epithelialization.

Mitotic cells can frequently be seen abutting rosettes or as part of the rosette in crosssections in confocal microscopy and EM (Fig. 1T; Fig. 2D,E,F). In addition, mitotic cells are often detectable at the next stage of MET, apically in the Co-Ep before separation of the epithelial layers (Fig. 1S,T; 2D). A three-dimensional analysis of mitotic cells in whole mount view revealed a 28% elevation in the frequency of mitotic cells in the transition zone as compared to the mesenchymal and epithelial zones (Fig. 3K). Since cell-cell and cellmatrix interactions are altered as cells round up upon entering mitosis and then reform upon mitotic exit^41–43^, it is possible that mitosis plays a role in resolving the rosettes into a mature epithelium by loosening intra-rosette cell-cell interactions and permitting post-mitotic rosette cells to form new junctions with neighboring lateral epithelial cells (see Discussion).

### Fibronectin Dynamics during MET

As described above, the first sign of incipient apical-basal polarity in the LPM is the accumulation of FN along the future basal surface of the Co-Ep in a lateral-to-medial fashion (Fig. 1I-L). This observation, together with published results implicating basally-located ECM in orienting epithelial polarity in other contexts^28^, led us to investigate whether ECM cues play a role in MET within the LPM.

First, we were interested to understand the dynamics of FN accumulation on the basal side of the LPM. mRNA expression analysis revealed that FN is mainly expressed in the ectoderm and weakly in the anterior endoderm, but is undetectable in the mesoderm (Fig. 4A). Thus, future basal FN must be recruited and assembled from non-mesodermal sources, as also occurs in somites^44,45^. Electroporating GFP-tagged FN into the primitive streak (thereby labeling primarily mesodermal cells) labeled all native FN-containing structures in both the mesoderm and ectoderm (Fig. 4B-B’’). This indicates that these structures are highly dynamic and constantly incorporating newly-secreted FN from the surrounding environment, and suggests that there is tight regulation to preserve the FN pattern regardless of the FN source. The alpha chain of the main embryonic receptor for FN in the chick, integrin α5β1 (Intα5) is expressed in a reciprocal manner to FN, i.e. mostly in the mesoderm, but not in the ectoderm (Sup. Fig. S3D-F)^44^, while integrin β1 (Int-β1) is expressed in all germ layers (Sup. Fig. S3G-I).

**Figure 4.**
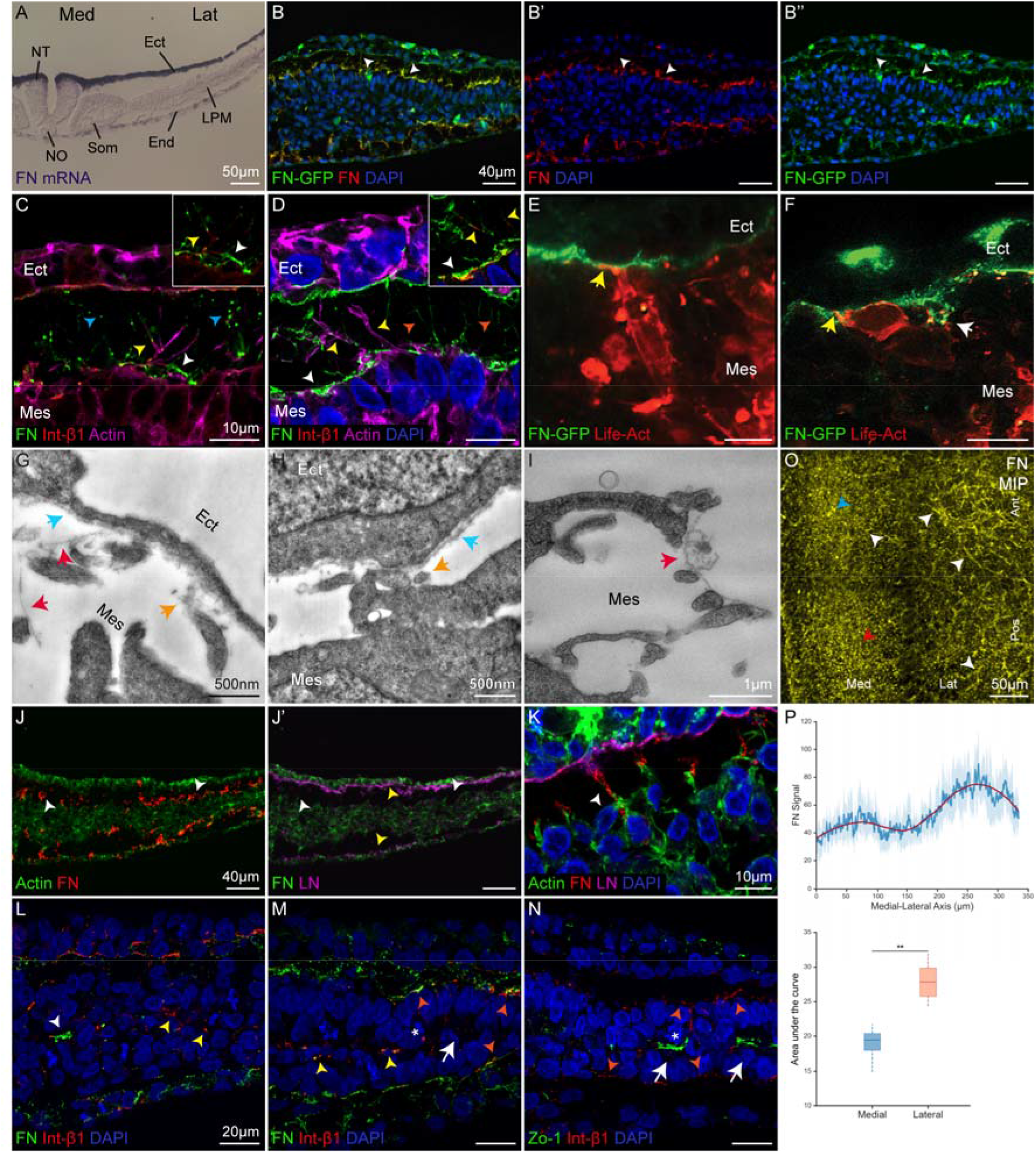
Cell-matrix interactions during MET. Medial is on the left. (A) In situ hybridization for FN, showing strong expression in ectoderm, weak expression in endoderm, and undetectable expression in mesoderm. (B) Section of embryo electroporated with full-length GFP-tagged FN and stained with an antibody to FN that recognizes both endogenous and electroporated FN. All of the FN on the basal LPM surface exhibits double staining with FN and GFP antibodies. (C,D) Consecutive stages showing FN secretion as punctate aggregates (blue arrowheads) that bind to Int-β1-coated filopodia (yellow arrowheads), and the subsequent formation of fibrillar extensions of FN at the ectoderm-mesoderm interface (orange arrowheads). White arrowheads point to basal FN-Int-β1 accumulation. (E,F) Embryo co-electroporated with Life-Act Red in the mesoderm and FN-GFP in the ectoderm during early stages of the MET process. (E) Shows strong interactions between mesodermal cellular extensions and the ectodermal basement membrane (BM) (yellow arrow). (F) Shows a mesodermal cell lying on the ectodermal BM and extending filopodia towards it (yellow arrow). White arrow in F indicates punctate FN secreted from the ectoderm binding to mesodermal cells. (G-I) Electron microscopy images showing interactions of mesodermal cellular extensions (G) and mesodermal cells (H) with ectodermal and mesodermal (I) ECM. Blue arrows indicate ectodermal BM. Red arrows indicate fibrillar ECM structures. Orange arrows indicate ECM aggregates. (J,J’) FN but not LN is detectable on the basal side of the LPM during the period of MET (white arrowheads), while LN is strongly expressed in the ectodermal and endodermal basement membranes (yellow arrowheads). (K) Mesodermal filopodia associated with FN fibrils that contact the ectodermal LN-containing ectodermal BM. (L-N) Confocal images from three stages of MET showing relocalization of Int-β1from a broadly-distributed pattern in the mesenchymal zone (L,M yellow arrowheads) to a basal pattern in the MET rosette-transition zone and epithelium (M,N orange arrowheads). White arrows indicate rosettes. White arrowhead in L indicates intramesodermal FN-Int-β1 aggregate. (O) Accumulation of FN during MET. Maximal Intensity Projection (MIP) of confocal Z-series of a flat-mounted FN-stained embryo showing transition from a punctate pattern (red arrowhead) containing aggregates (blue arrowhead) to a fibrillar pattern (white arrowheads). Note that the latter form is more abundant laterally, especially anteriorly. (P) Top panel shows line plot of averages of FN intensity on the medial-to-lateral axis, and bottom panel compares sum of FN intensity in the medial and lateral halves of the LPM undergoing MET of a representative embryo.

High resolution confocal imaging of immunostained embryos for actin, FN and Int-β1, and embryos electroporated with FN-GFP and Life-Act-mCherry, revealed numerous FN-associated Int-β1-coated filopodia extending between the nascent Co-Ep and the overlying ectoderm (Fig. 4C-F). Electron microscope imaging revealed a close interaction between filopodia and extracellular matrix structures present at the ectoderm-mesoderm interface and the ectodermal BM (Fig. 4G-I). Taken together, these findings suggest that integrin-coated actin-filopodia may play a role in shuttling FN to the future basal side of the Co-Ep and/or transmitting an integrin-mediated signal. These filopodia also interact with the LN-containing ectodermal BM (Fig. 4K), although the LPM itself does not accumulate LN until well after epithelialization (Fig. 4J’; Sup. Fig. S1H,K,J).

Int-β1, similar to the apical markers, is expressed in a dispersed pattern in mesenchymal cells (Fig. 4L,M, yellow arrows), sometimes colocalizing with the intra-mesodermal FN aggregates that are visible initially (Fig. 4L, white arrow). Subsequently, Int-β1 is translocated from the apical sides of the LPM to the forming FN layer basally (Fig. 4L-N). Notably, FN and Int-β1 are strongly expressed on the basal side of the rosette structures (Fig. 4M, orange arrows).

As epithelialization proceeds, FN undergoes an increase in expression level and in apparent complexity of the fibrillar network (Fig. 4O-P). However formation of FN fibrils does not appear to be required for MET, as electroporation of a 70kD FN fragment that acts as a dominant negative to prevent FN fibril formation^46^ did not prevent MET (Sup. Fig. S4). This suggests that if FN does play a role in MET, it may not need to be in fibrillar form.

### The Integrin Signaling Axis is Required for MET

Upon secretion, FN dimers bind to integrin receptors on the cell surface, which interact with cytoskeletal adaptors including talin and activate downstream signaling targets including focal adhesion kinase (FAK), thus forming focal adhesion complexes important for rearrangement of the cytoskeleton and intracellular signaling^47–50^. In addition, through “inside-out signaling”, cellular tension initiated by FN binding to integrin receptors promotes assembly of FN fibers in the ECM^51^. We conducted experiments to investigate the role of the ECM-Integrin axis in MET.

Immunostaining for phosphorylated FAK (pFAK) revealed that it is expressed initially weakly throughout the mesoderm (Sup. Fig. S5A,A’, red arrow), and relocalizes basally (Sup. Fig. S5A,A’,B,B’, white arrows), and becomes highly upregulated as epithelialization proceeds (Sup. Fig. S5C-C’’). In order to functionally assess a possible role for FAK signaling during MET, we electroporated a dominant negative GFP-labeled FAK construct (FF-GFP)^52^ into the future LPM cells (N=24 embryos in 5 experiments). Note that electroporation introduces the construct into a subset of the cells, so that the LPM becomes a mosaic of electroporated and non-electroporated cells. We found that heavily electroporated areas exhibit large masses of FN-demarcated unorganized cells (Sup. Fig. S6A-D,I-L). In areas with lower numbers of electroporated cells, the FF-GFP electroporated cells lie flat along the basal side, lacking dorso-ventral elongation, and do not integrate into the epithelial layer, which is constituted mostly by wild-type cells (Fig. 5A-C, white arrows). This suggests that cells with defective FAK signaling are excluded from participating in MET and extruded from the forming epithelial layers. Notably, the basally-extruded cells exhibit extensive cellular processes and are embedded in the underlying FN-containing ECM (Fig. 5G-G’’), indicating that they retain mesenchymal properties. Consistent with this interpretation, Actin, Zo-1, aPKC and Int-β1 staining in FF-GFP electroporated cells resembles that of the mesenchymal state, with multiple Zo-1 and Int-β1 dispersed punctae distributed throughout the cell (Fig. 5A’,C’), while aPKC staining is absent (Fig. 5B,B’). If sufficient nonelectroporated cells remain in the mesodermal layer following the basal exclusion of FF-GFP-electroporated cells, the non-electroporated cells can still form a seemingly normal epithelial layer (Fig. 5C; Sup. Fig. S6Q-T). In anterior regions, extruded electroporated cells may clog the opening of the coelomic cavity (Sup. Fig. S6Q-T). In summary, FF-GFP hinders the MET process and prevents the apical-basal polarization of the electroporated cells. In the absence of FAK signaling, cells maintain their mesenchymal properties, are unable to integrate into the forming epithelial layers, and are extruded either basally or apically.

**Figure 5.**
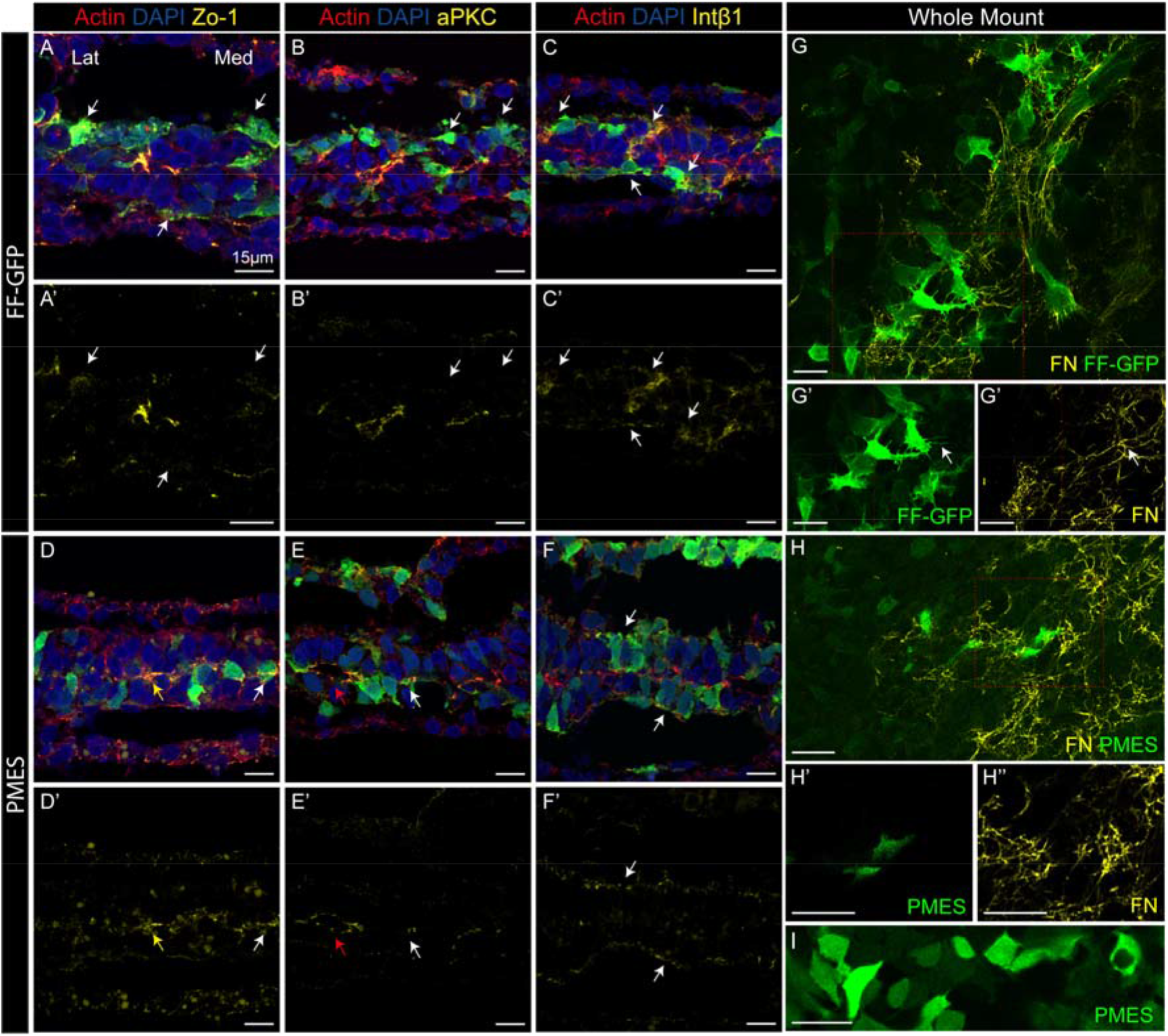
Effects of interference with FAK function on MET. Medial is on the right. (A-F) Sections of embryos electroporated with the dominant negative FAK construct FF-GFP (AC) or control construct PMES (D-F). Note basal exclusion of FF-GFP-electroporated cells from the epithelial layer (white arrows), while PMES-electroporated cells are integrated into the epithelium. FF-GFP-electroporated cells also exhibit a mesenchymal pattern of the apical markers Zo-1 (A’) and aPKC (B’), and intracellular accumulation of the basal marker Int-β1 (C’). Yellow arrows indicate rosette. Red arrows indicate coelom. (G-I) FF-GFP-electroporated (G) or control-electroporated (H,I) cells imaged at the basal surface (G,H) or middle (I) of the LPM. There are abundant FF-GFP-electroporated cells basally and they exhibit many cellular processes that are embedded in the basal FN network (G-G’’, arrow) as compared with very few control cells (H), which exhibit many fewer processes (I).

Talin is a component of focal adhesions that links the ECM and integrin to the cellular cytoskeleton and has been previously reported to be involved in integrin regulation and activation^53^. Electroporation of a dominant negative talin construct (N=13 embryos in 3 experiments) consisting mainly of the head domain (qTalinH)^54^, caused a more considerable disruption in epithelialization compared to FF-GFP (Fig. 6A,B,G; Sup. Fig. S7A-C,I-K; compare with Fig. 6D,E,H; Sup. Fig. S7E-G,L-N, and Fig. 5). Similar to FF-GFP, qTalinH-electroporated embryos exhibit clumped masses of cells (Fig. 6A,B,G) that are often demarcated by FN (Sup. Fig. S7A-D). In addition, these embryos are characterized by a dispersed, mesenchymal pattern of apical markers N-Cad, Zo-1 and aPKC (Fig, 6B’,C,G’; compare with E’,F,H’). In contrast to the FF-GFP phenotype, however, the effects of qTalinH were not exclusively cell-autonomous. Surrounding non-electroporated cells were also often defective in MET progression (Fig. 6B,B’,C,G,G’; Sup. Fig. S7A-D,I-K). Consistent with this, whole mount imaging revealed an expansion of the transition zone and line of rosettes (Fig. 6K), defects in development of the epithelial hexagonal N-Cad pattern (Fig. 6I), and depletion of N-Cad in more lateral regions since most of the cells are aggregating more medially (Fig. 6I,K,O). Although FN basal expression was slightly less dense in cross sections (Sup. Fig. S7D), in whole mount the FN network was strikingly affected, exhibiting a sparse pattern (Fig. 6M), when compared with control (Fig. 6N) or FF-GFP electroporated (Fig. 5 and Sup. Fig. S6) embryos, and in line with previous reports of the importance of talin to integrin activation^53^. Together, these observations suggest that talin inhibition prevents the cells from progressing in the MET process, thus remaining in the primitive mesenchymal and transitional states.

**Figure 6.**
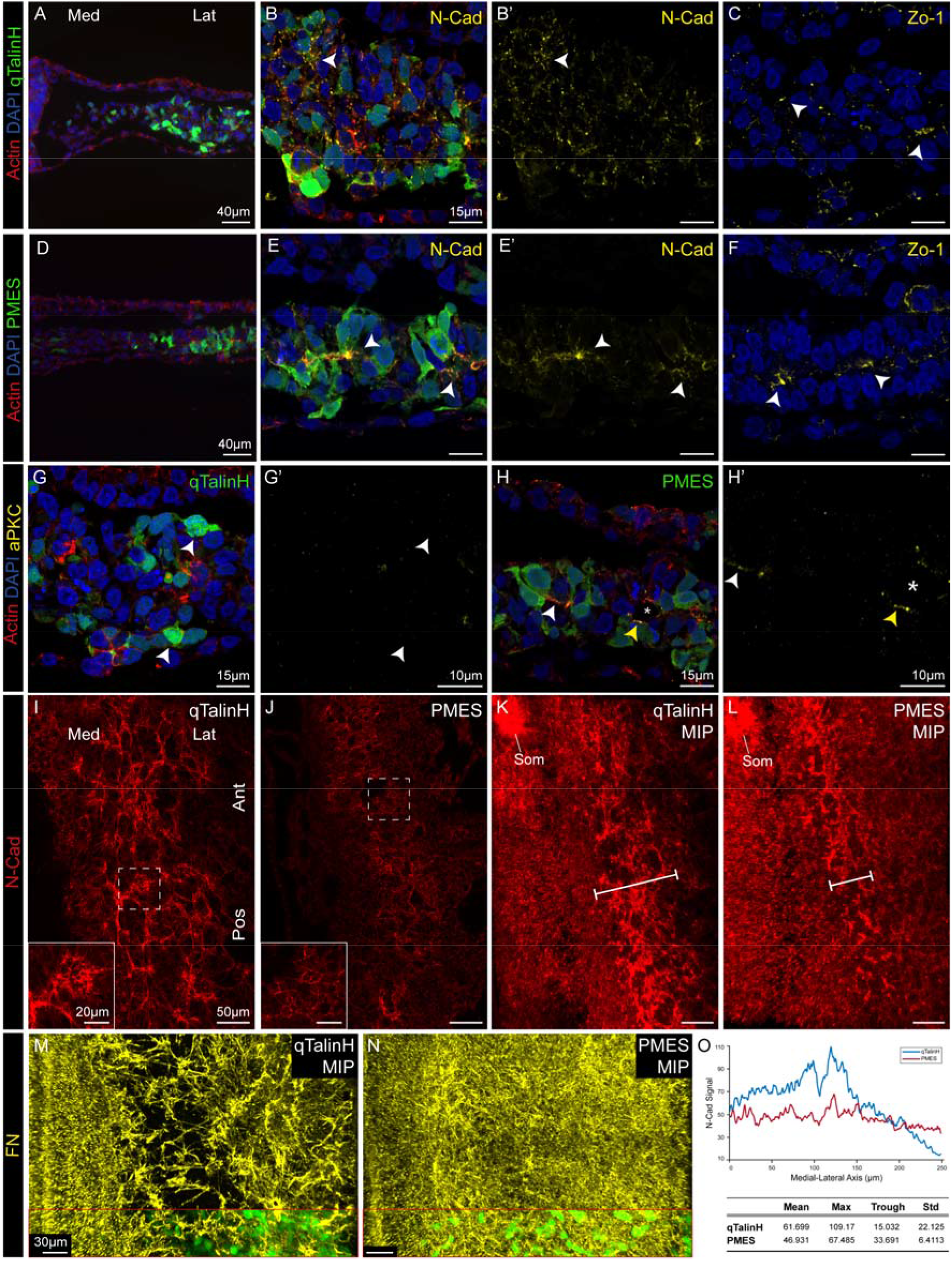
Talin inhibition disrupts MET progression. Medial is on the left. (A-H) Sections of embryos electroporated with dominant negative Talin construct qTalinH (A-C,G,G’) or control plasmid (D-F,H,H’). A and D are widefield and B,C,E,F,G,H are confocal views. qTalinH-electroporated embryos maintain a mesenchymal phenotype, both morphologically and with respect to distribution of polarity markers in both electroporated and nonelectroporated cells. Note the thickness of the LPM in qTalinH-electroporated embryos (B,G). (I-N) Whole mount views showing the coarser staining patterns of N-Cad (I) and the wider “line of rosettes” (K) and sparser FN-network (M) in qTalinH-electroporated embryos. The insets in M and N show GFP in a representative area of each embryo to document the widespread GFP distribution. (O) Quantification from Maximal Intensity Projections (MIP) of N-Cad signal along the X axis in qTalinH vs. control embryos. Note the largely uniform signal intensity in PMES-embryos compared to the high peak and low trough in qTalinH-embryos. Som, somites.

In summary, these results indicate that integrin-mediated signaling via FAK and Talin is prerequisite for MET.

## Discussion

Based on the current work we propose a model for MET in the lateral plate mesoderm (Fig. 7). Before MET is initiated, the LPM is multilayered and comprised of cells that are loosely connected, mesenchymal in morphology i.e. lack polarity, and extend filopodia that interact with the ECM and neighboring cells (Fig. 7A, left side). However, these cells do express many proteins associated with cell polarity, including N-Cad, Zo-1, and Par-3, but not aPKC. Although fairly colocalized, these markers are dispersed throughout the cell membrane in a non-polarized manner. Subsequently, ECM accumulation on the future basal side of the epithelium, acting through integrin and its downstream effectors FAK and Talin (Fig. 7A, right side), promotes cell intercalation and provides a polarizing signal that redirects these apical components to the incipient apical sides of the cells concurrently with the emergence of aPKC apically. The formation of actomyosin contractile rings produces apically-constricted trapezoidal cells that are brought together by adherens and tight junctions to form rosette-like structures. Finally, possibly aided by the remodeling of cell-cell junctions that occurs during mitosis, polarized cells of the rosettes re-orient their cell-cell junctions to become incorporated into the adjacent mature double-layered epithelium.

**Figure 7.**
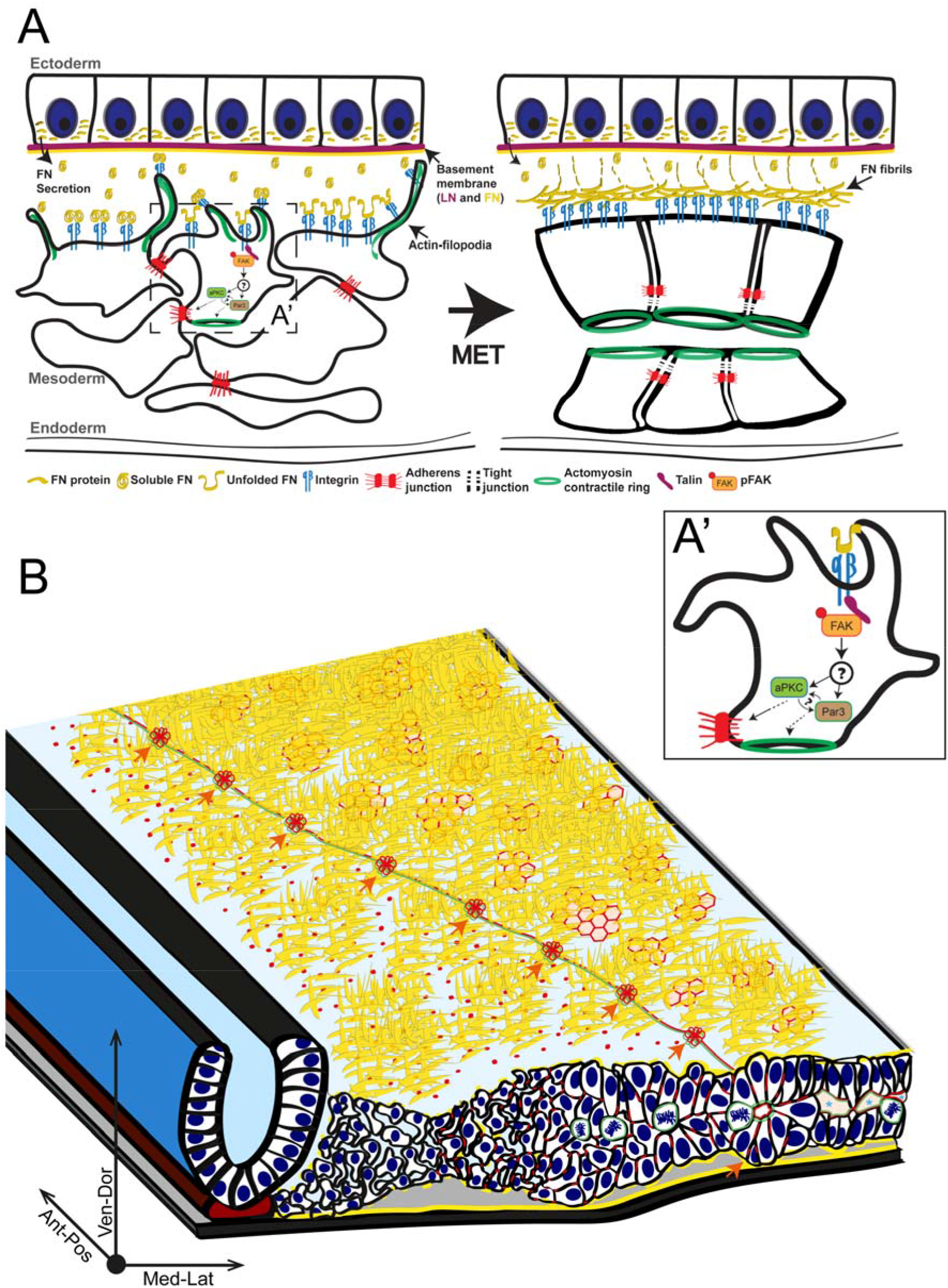
Proposed Model of MET in the LPM. (A) Cross-sectional view of mesenchymal and epithelial stages of MET. Note the changes in FN network, cellular morphology including filopodia, and junctional distribution. (B) Whole-embryo view showing the diagonal line of rosettes as seen in a flat whole mount (XY plane, from above) and an appropriate cross-section (XZ plane) depicting the stages of MET in detail. In the XY plane, note the increase in the richness of the FN network in the lateral and anterior directions (yellow strands) and the change in N-Cad expression from dotted-to-hexagonal on moving from the medial to the lateral side (depicted in red). In the XZ plane, note the increase in the density and decrease in thickness towards the lateral side, the changes in the junctional complexes, the rosette structure, the location of mitotic cells, and opening of the coelom. Asterisk indicates coelom. Orange arrows indicate the line of rosettes. Actin is depicted in Green. See text for further discussion.

The current work reveals several novel characteristics of MET in the lateral plate mesoderm and also sheds light on the MET process more generally. First, we observed that MET occurs as an organized wave, sweeping diagonally from anterior and lateral to posterior and medial, thereby producing the line of rosettes that extends in a complementary direction from anterior-medial to posterior-lateral. Several possible, non-exclusive mechanisms can be postulated for propagation of the wave. One is propagation via the ECM, in which mature epithelium causes a corresponding change in the ECM that is propagated locally to induce MET in neighboring cells. We found that inhibition of FN fibril formation does not affect MET (Sup. Fig. S4), but other changes in the ECM remain a possibility for wave propagation, such as the recruitment of FN dimers or other ECM components. A second possibility is a signaling event in which a factor secreted by already-epithelialized cells induces MET in their more medial neighbors. Such a mechanism occurs during kidney tubule formation, in which Wnt signaling from the epithelial nephric duct initiates aggregation and epithelialization in the neighboring metanephric mesenchyme^6,30^. Although BMP7 expression in the ectoderm has been reported to be required for coelom formation^55^, this factor in itself would not be able to account for the wave-like nature of MET in the LPM, since BMP7 is expressed uniformly throughout the ectoderm^56^. A third possibility is a cell autonomous mechanism. A characteristic feature of the MET wave is that it proceeds from anterior-lateral to posterior-medial. Interestingly, fate maps have shown that, after gastrulation, LPM cells migrate in an anterior-lateral direction^35,36^. Thus anterior-lateral cells are the embryologically “oldest” cells of the LPM (they have spent the longest time as mesoderm since their gastrulation). It has been found that somite formation (another example of MET) requires reduction in levels of FGF signaling. FGF mRNA stops being expressed in presomitic cells when they gastrulate, and the natural decay in levels of FGF mRNA and protein serves as a molecular clock to regulate the timing of the onset of somite formation^57^. Analogously, LPM cells may be expressing an inhibitor to epithelialization that must decay before MET can be initiated, explaining why the “oldest” cells initiate MET. Finally, there could be a long-range secreted inhibitor of MET that is located posterior-medially in the embryo, such that MET can only occur once cells are sufficiently distant from the inhibitor.

The existence of the morphogenetic wave aided in understanding the stages of MET, since at any one time the stages are arranged spatially within the embryo. This regular spatial arrangement facilitated the identification of another characteristic of MET in the LPM: the formation of rosettes as a transitional structure between mesenchymal and epithelial forms. A structure termed “rosette” has been described during epithelial remodeling in the mouse egg cylinder^58^, Drosophila germ band extension^59^, and other systems^60^, but has been regarded as a transient epithelial structure that mediates cell rearrangements within the epithelial layer. The current observations present rosette-like structures as a stage during the epithelialization process itself, and also uncover several novel characteristics of the rosette. First, while previous studies have described rosettes as discrete structures, the current studies reveal that, at least in the case of the LPM, they have a complex three-dimensional morphology, as seen in 3D reconstructed whole mounts and the supplemental videos. Furthermore, what is unique in the LPM is the formation of a line interconnecting the polarized cell aggregates, which we term the “line of rosettes”, which represents the transition zone between mesenchymal cells on one side of the line and epithelial cells on the other side and which sweeps diagonally over the LPM. The line of rosettes is normally quite thin in width, comprising only one or a few aggregates, indicating that the transition from mesenchyme to epithelium occurs quite rapidly.

An important aspect of rosette biology is their resolution into a mature, two-layered epithelium. Cells of the rosette have apical junctional complexes including tight junctions and adherens junctions and are thus closely attached to one another. In order to form a mature epithelium, some of these cells must break intra-rosette cell-cell junctions and form new ones with neighboring cells of the mature epithelium. It was intriguing to observe an enrichment of mitotic figures in the vicinity of the line of rosettes (Fig. 2D’,E,F; 3J; 4M; 7B orange arrow). During mitosis, epithelial cells round up which decreases their apical tension, loosen their attachment to their neighbors and with the basement membrane, and reform them again after cell division is complete^41–43^. It is interesting to speculate that mitosis may play a role in weakening cell-cell connections within the rosette. Subsequently, the neighboring epithelial cells could “entrain” former rosette cells and induce them to join the mature epithelium. Mitosis has been previously suggested to play a role in morphogenesis and cell-mixing within an epithelium^43,61–63^, but has not, to our knowledge, been implicated in MET. Antagonization of apical constriction may also play a role in rosette resolution independent of mitosis. In the future it will be important to understand the basis for the mitotic enrichment and whether it results from growth factor signaling, a prolonged mitotic phase, or other cause.

Interestingly, mitotic cells were also frequently observed at the last stage of MET, prior to coelom formation (Fig. 2D’ red arrow; 4N asterisk; also depicted in the model Fig. 7B, the most lateral side). Therefore, we postulate that breaking cell-cell adhesions during mitosis might aid in the opening of the coelomic slits. Alternatively or cooperatively, the negatively-charged sialomucin podocalyxin that emerges late during MET and coats the apical surfaces might also play a role in the separation of the two epithelial layers. Podocalyxin has been implicated in the separation of podocyte foot processes^39^, and in lumenogenesis in the aorta, MDCK cells, and the mouse epiblast.^28,64,65^

While most apical proteins (including N-Cad and Zo-1) were expressed at similar levels in mesenchymal and epithelial cells, Par-3 showed a modest but steady rise during epithelialization, and aPKC exhibited a dramatic jump in expression between the mesenchymal and the transition (rosette) stages. aPKC mRNA did not exhibit the same jump, indicating that the change in protein levels was post-transcriptional, either due to increased protein translation or stability, or a conformational change that could be identified by the antibody used in this study. It is interesting to consider that N-Cad and Zo-1 are structural components of the apical junctional complex, whereas Par-3 and aPKC play scaffolding and regulatory roles^24,25,66,67^. aPKC has been found to be critical for establishment and maintenance of apical-basal polarity in other systems^68,69^, and Par-3 has been found to be required for proper localization of aPKC to the apical-lateral domain interface^25,69,70^. The current findings suggest that accumulation of aPKC may play a key role in progression of MET within the LPM.

The current study presents evidence that ECM-integrin signaling, acting through FAK and Talin, plays a critical role in the initiation and propagation of MET in the LPM. In the model presented here, ECM accumulation at the future basal side of the epithelium activates integrin signaling and provides a positional cue to initiate apical-basal polarity. Many components of the polarity apparatus and integrin are already expressed in mesenchymal cells, so it appears that the ECM signal primarily triggers reorganization of these components to their destined domains. Knockout mice for Int-β1 have impaired polarization and epithelialization in diverse tissues^71^. Integrin signaling has also been found to be important for epithelialization of cells in the mouse egg cylinder and cultured MDCK cells^28,29^. In the case of the egg cylinder, the ECM cue is thought to be laminin (LN). However, in the LPM, LN starts accumulating basally only after epithelialization is already well underway (Sup. Fig. S1). In contrast, FN accumulation on the basal side of the forming epithelia is the earliest indication of epithelialization of the LPM, suggesting that FN may promote epithelialization. The prominent filopodia observed in LPM cells at the mesenchymal stage may be a mechanism to recruit FN that is secreted from the neighboring ectoderm. Alternatively, these filopodia may also interact with the LN-containing basement membrane of the ectoderm, which may provide a basal polarizing cue, or with other secreted molecules at the mesoderm-ectoderm interface.

Inhibition of integrin signaling in our experiments (FF-GFP and especially qTalinH) led to the formation of multilayered large masses of cells that were unable to advance in the epithelialization process, with an expansion of the transition zone in qTalinH-electroporated embryos and a defective FN network. Int-β1-deficient ESCs aggregate but cannot polarize in culture^28^ and ECM-integrin signaling and the FN matrix have been reported to be essential for intercalation and convergent extension in Xenopus^72^. In addition, rosette-like structures have been proposed to mediate intercalation necessary for convergent extension in Drosophila germ band extension, in vertebrate kidney tubule and primitive streak morphogenesis, and in neural tube closure.^59,60,73,74^ Since FN accumulation precedes cell intercalation and rosette formation and there is always a rich FN network outlining the rosettes in the LPM (Fig. 4M,N), it is intriguing to suggest that ECM-integrin signaling guides the gross morphological remodeling occurring in the LPM from multilayered mesenchyme to rosette formation to epithelial sheets, through the described molecular and structural intermediate stages.

All major regions of the mesoderm undergo MET. The paraxial mesoderm forms somites, the intermediate mesoderm (IM) forms kidney tubules, and the LPM forms the coelomic epithelium. These epithelia have different morphologies. In particular, the somite and kidney tubules remain discrete epithelial structures. In contrast, in the LPM, the rosettes act as epithelial-like transient structures that resolve into two, huge epithelial sheets. Thus, while there are shared features between these cases of MET, there are also important differences. It is interesting to wonder why, for example, the IM forms multiple small epithelial aggregates that remain discrete and do not fuse with each other, such that each will form a separate kidney tubule, while in the LPM there is rosette formation and sequential resolution to enable integration into cohesive epithelial sheets. The current findings suggest that the difference may lie in the dynamic nature of the apical surface and the junctional complexes in the LPM. In contrast, cell-cell connections in nascent kidney tubules or somites may be stronger and less susceptible to remodeling. Thus, through varying strength of cellcell junctions, different tissues may generate different epithelial morphologies.

## Acknowledgements

We thank Masatoshi Takeichi, Yuki Sato, Ben Allen, Paris Skourides, and Kelly McNagny for providing plasmid and antibody reagents. Ariel Shemesh assisted with image processing and quantification. We also thank Maya Holdengraber and Melia Guruwitz from the Interdepartmental Imaging Facility for assistance with imaging. We thank Jad Asleh for assistance with data analysis. This work was supported by grants to TMS from the Israel Cancer Research Fund, the Israel Science Foundation (2407/15), and the Rappaport Family Foundation. M.A. was supported by a scholarship from the Clore Israel Foundation.

## Methods

### Chicken Embryos

Studies were carried out on fertilized Hyline strain chicken eggs (Moshav Orot, Israel) and incubated in the lab at 38.6 degrees C in a humidified incubator until the desired stage.

### Plasmids

PMES drives constitutive expression of an inserted gene and co-expresses eGFP under an IRES element^75^. Focal adhesion kinase (FAK) dominant negative construct (FF-GFP)^52^ was kindly provided by Paris A. Skourides and cloned into the PMES vector. Dominant negative quail Talin head construct (qTalinH), FN-GFP, and FN 70kD were generously provided by Yuki Sato^54^. Life-Act mCherry was purchased from Addgene (#54491) and cloned into pCAG^76^ to drive expression in the chick embryo.

### Electroporation

HH stage 4 chick embryos were harvested by attaching them to a paper ring and placed dorsal-side up on the platform of an electroporation chamber over the positive base-electrode. For primitive streak electroporation, DNA solution mixed with Fast Green dye was injected into the posterior half of the primitive streak. Next, the negative upper electrode was positioned superficially in the liquid and used to deliver 6 pulses of 9 volts.

For ectoderm electroporation, the needle was inserted superficially underneath the vitelline membrane, on one of the lateral sides, starting anteriorly and slowly moving posteriorly whilst injecting the DNA. Afterwards, the negative upper electrode was positioned superficially in the liquid, initially more anteriorly, and then gradually more posteriorly in the same pattern in order to label a wide area of the ectoderm. Fluorophore expression can be seen in the LPM or the ectoderm within 3 hours of electroporation. After electroporation, the embryos were incubated on agar-albumin dishes ventral-side up in 38°C for 16-24 hours (until they reached HH stage 10).

### Immunofluorescence on Sections

Embryos were fixed in 4% paraformaldehyde, embedded in 7.5%gelatin/15%sucrose and cryosectioned at 10 or 20 μm. For, Zo-1, pFAK, and Intβ1 embryos were fixed for 1h at RT. For Par-3 fixation was for 1h at 4 degrees. For other markers fixation was overnight at 4 degrees. Immunofluorescence staining was performed with the following primary antibodies: rabbit anti-GFP (1:1000, Molecular Probes a-6455), mouse anti-GFP (1:500, Molecular Probes3E6), rat anti-RFP (1:1000, Chromotek), mouse anti-LN (1:10, DHSB clone 3H11), rabbit anti-LN (1:300, Sigma L9393), rabbit anti-Zo-1 (1:1500, Invitrogen 40-2200), mouse anti-N-Cad (1:5, DSHB clone 6B3), rabbit anti-FN (1:400, Sigma F3648), mouse anti-FN (1:30, DSHB clone B3/D6), mouse anti-Podocalyxin (1:500, a gift from K. McNagny), mouse anti-Intβ1 (1:10, DSHB clone V(2)E(9)), rabbit anti-Par-3 (1:500, Millipore), mouse anti-aPKC (1:500, Santa Cruz), rat anti-RFP (1:1000, Chromotek), mouse anti-Arl13B (1:1000, NeuroMAB) and rabbit anti-pHistone3 (1:500, Millipore), and mouse anti-pFAK (1:200, ThermoFisher). Actin filaments were stained with ActinGreen488 (1:25, Invitrogen R37110) or ActinRed555 (1:25, Invitrogen R37112). Nuclei were labeled with DAPI (Sigma) at a concentration of 1μg. The secondary antibodies that were used (all from Jackson Immunoresearch) include Cy3 Donkey anti-mouse IgG (1:250), Cy3 Donkey anti-rabbit IgG (1:250), Cy3 Donkey rat IgG (1:250), Alexa Flour 488 Donkey anti-rabbit F(ab) IgG (1:250), Alexa Flour 488 Donkey anti-mouse F(ab) IgG (1:250), Alexa Fluor 647 Donkey anti-mouse IgG (H+L) (1:250), Alexa Fluor 647 Donkey anti-rabbit IgG F(ab) (1:250), and Alexa Fluor 647 Donkey anti-rabbit IgG (H+L) (1:250). Immunostaining was performed as described in Nishimura et. al 2008^73^. Immunofluorescence-labeled sections were mounted using Fluorescence Mounting Medium (DAKO S302380) and imaged with either a Zeiss Axioimager M1 widefield epifluorescence microscope and a Qimaging ExiBlue monochrome camera, or a Zeiss LSM700 confocal microscope. Image overlays were performed with ImagePro Plus software (Mediacy) or Imaris software.

### Whole Mount Immunostaining and Clearing

Whole embryo immunofluorescence was performed using the same antibody concentrations and fixation conditions as for cryosections, except for N-Cad, which was used at 1:20 with Alexa Fluor 647 Donkey anti-mouse IgG, and 1:100 with Cy3 Donkey anti-mouse IgG, and Int-β1, which was used at 1:50. For immunostaining, the embryos were incubated with the primary antibodies and ActinGreen or ActinRed for 48h at 4 degrees, washed with triton 0.3%, and then transferred to secondary antibodies and DAPI for 48-72h. Subsequently, for clearing, they were washed and then treated with increasing concentrations of fructose from 25% to 100% over a span of three days. Finally, they were transferred to SeeDB solution^77^ for one night. On the following day, the embryos were placed with SeeDB solution in a double-sided-tape-generated rectangular well on a slide and enclosed with a coverslip.

Imaging was performed with Zeiss LSM880 confocal microscope using 25x/NA0.8 and 63x/NA1.1 PlanApo long working distance objectives. Zen software was used for stitching, deconvolution, and analysis. Imaris software was used for stitching, three-dimensional (3D) view, maximal intensity projection (MIP), and ROI 3D-segmentation and clips. CellProfiler was used for colocalization analysis of at least 15 different Z-levels for each pair.

### Vibratome

Embryos were embedded in 4% agarose, sectioned using a vibratome at 100-150 μm. Immunostaining and clearing were done similar to whole embryos. The sections were placed in a well, covered and imaged with either Zeiss LSM700 or Zeiss LSM880 confocal microscopes.

### In-situ hybridization

Whole mount in-situ hybridization (ISH) was performed as previously described^78^ using digoxigenin-labeled RNA probes for FN, Intβ5, Intβ1 and PKCζ using the following sense oligos: For FN, GAGAGC AAACCACCAAGCTC and CTTTCTCCTGCC GCAACTAC; for, Intβ5 AGGTGCTGAGGGGGCAA and CACGACGGTGAGCGAAG; for Intβ1 CCAAAGAAAGGGCAAAATGA and ACGCTAAATGGCTAGGCAGA; and for PKCζ CCAGAAGATGGAGGAAGCTG and ATACACCAAGAGCCCACCAG. Intβ5 probe was kindly provided by Solveig Thorsteinsdottir^44^. Whole embryos were photographed with a Leica MZ16FA stereomicroscope and Leica 420C camera. Subsequently, the embryos were cryosectioned at 20μm and imaged using a Zeiss Axioimager M1 widefield microscope with DIC optics and a Qimaging ExiBlue monochrome camera with RGB color filter.

### Electron Microscopy

Embryos were fixed in 3mm plates with 2% glutaraldehyde and 2% paraformaldehyde in 0.1 M Sodium Cacodylate buffer pH 7.4 containing 5 mM CaCl2 for 2 hours at Room Temperature (RT), then cleaned and moved to glass vial with fresh fixation solution for 30 minutes at RT. Following fixation, embryos were washed with Sodium Cacodylate buffer, post-fixed and stained for 1 hour on ice with 1% Osmium TetraOxide, 0.5% Potassium hexacyanoferrate, 0.5% Potassium dichromate in 0.1M Cacodylate buffer containing 2 mM CaCl2, following which the samples were en-block stained with 1% uranyl acetate for 1 hour on ice. Then the tissue was dehydrated in a graded ethanol series and embedded in Epon812. 75 nm ultrathin sections were cut with a ultramicrotome UC7 (Leica), transferred to copper grids and viewed using Zeiss Ultra-Plus FEG-SEM equipped with STEM detector at accelerating voltage of 30 kV or using Talos L120C Transmission Electron Microscope at accelerating voltage of 120 keV. For FEG-SEM, samples were coated with a thin layer of carbon to increase conductivity.

### Quantification and Statistical Analysis

#### Thickness

The thickness of posterior and anterior sections of ten different embryos was measured in ImageJ, at the same distance from the midline. Two-Sample T-Test was used to compare the means.

The thickness of lateral and medial regions of the mesoderm of the same sections of ten different embryos was measured in ImageJ. Paired-Sample T-Test was used to compare the means.

#### Density in 2D

Distribution and density of 15 mesenchymal and 7 epithelial cross sections were measured using Python libraries. Cell segmentation was performed using a pre-trained deep learning model. Manual segmentation was done on the epithelial sections to help further distinguish the abutting cells. Then, the data retrieved from segmented nuclei images were processed for calculating the density occupied by each cell group. Wilcoxon rank sum test was used to compare the means.

#### aPKC and Par-3 Intensity

Maximal intensity projection (MIP) of the confocal-imaged zstack was obtained using Imaris software. Signal intensity measurements were done on the MIP using ImageJ Plot Profile tool on fifteen parallel horizontal mediolateral lines (on the anterior-posterior axis). In Matlab, the mean and standard deviation of the data matrix were plotted using the shaded area error bar plot function^79^.

#### FN Intensity

The measurements were done similar to Par-3 and aPKC, with 8 parallel horizontal mediolateral lines (on the anterior-posterior axis). In Matlab, a Savitzky-Golay filter was used to smooth the data and then the approximate integral of the signal of each peak was computed via the trapezoidal method (“area under the curve”). Statistical analysis of the two areas was done using the Wilcoxon signed rank test.

#### N-Cad Intensity

The measurements were done similar to Par-3, aPKC and FN, but with 11 parallel diagonal lines perpendicular to the rosette line for both PMES and qTalinH embryos. In Matlab, the means of the lines were plotted and the mean, maximum, trough and standard deviation of each line were calculated.

#### Mitoses Analysis

HH stage 10 embryo immunostained for pHistone3, N-Cad, Actin and DAPI was imaged using a Zeiss LSM880 confocal microscope. Segmentation of the LPM region was performed using Imaris software. Image processing was performed on a Python platform (Anaconda) using image and data processing libraries, e.g., numpy, scipy, cv2, seaborn, skimage, etc. Each of the embryonic and mitotic nuclei 3D images were first preprocessed to improve their contrast, then 3D segmented using a pre-trained Convolutional Neural Network (CNN). Following segmentation, center of mass was calculated and plotted as iso-proportions of the nuclei density, i.e., each curve shows a level that some proportion of the density lies below it.

