## Supplemental Figures for "A Morphogenetic Wave that Generates Mesenchymal-to-Epithelial Transition in the Lateral Plate Mesoderm"

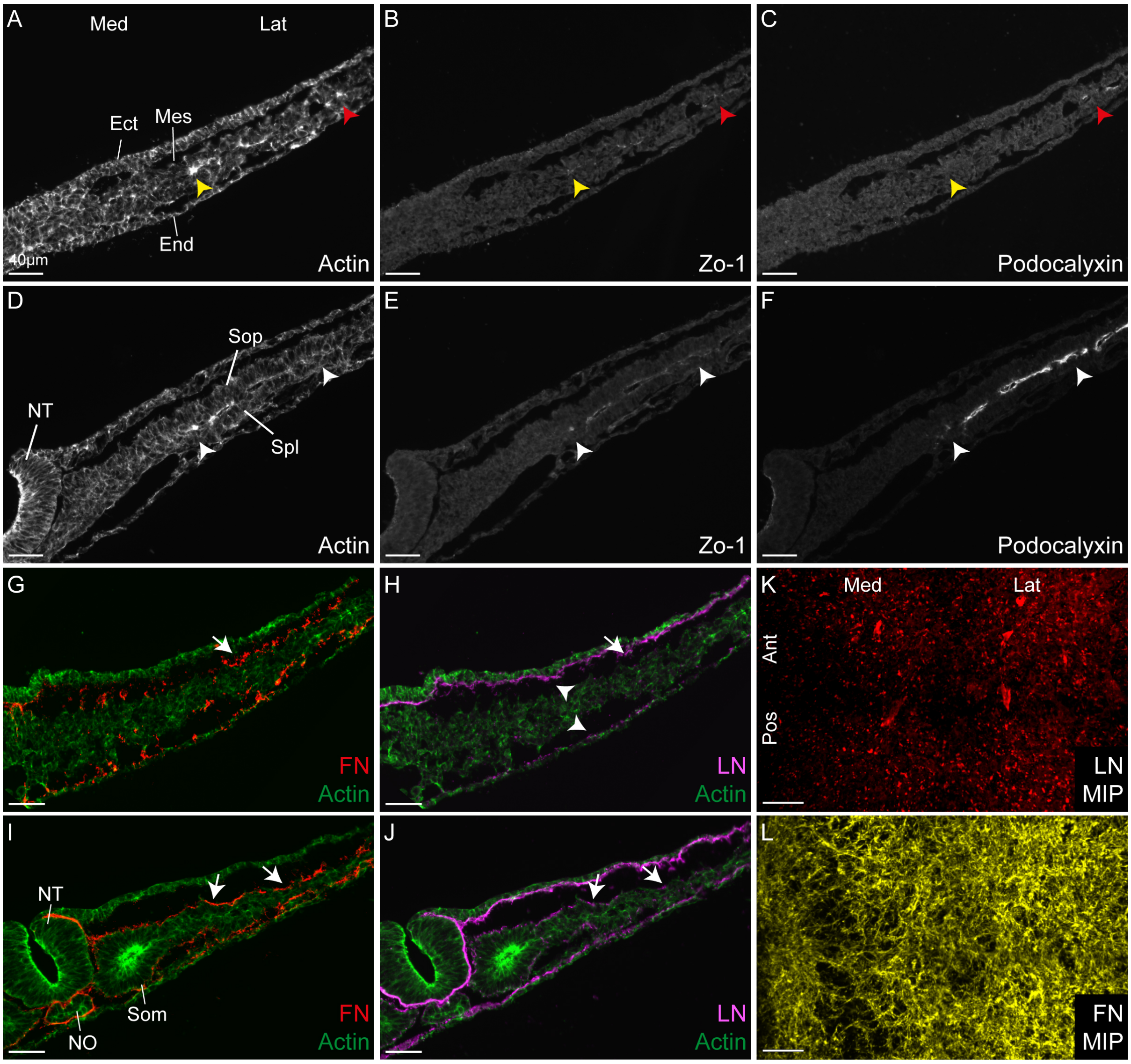


Supplemental Figure S1. (A-F) Podocalyxin is a late marker of MET. Early (A-C) and later (D-F) stages in the MET process. At early stages podocalyxin is visible only in the most lateral regions of the LPM (A-C red arrowheads), despite evidence for the onset of apical-basal polarization and epithelialization in more medial regions (A-C yellow arrowheads). At later stages (D-F), podocalyxin expression becomes aligned with other apical markers (white arrowheads). (G-L) Laminin accumulates late in the MET process. At early stages in MET (G,H), fibronectin (FN) has begun to accumulate on the basal side of the forming coelomic epithelium in lateral regions of the LPM (arrows), but laminin (LN) is not detected. At these stages LN is strongly detected in the basement membranes of the ectoderm and endoderm (arrowheads). At later stages in the MET process (I,J), LN is also detected on the basal side of the coelomic epithelium (arrows), though weaker than the LN of the ectoderm and endoderm. (K,L) Flat mount of HH stage 9 embryo stained for LN and FN. Note the punctate pattern of LN staining as opposed to the fibrillar pattern of FN. Ect, ectoderm; End, endoderm; Mes, mesoderm; NO, notochord; NT, neural tube; Som, somite; Spl, splanchnopleuric mesoderm; Sop, somatopleuric mesoderm.

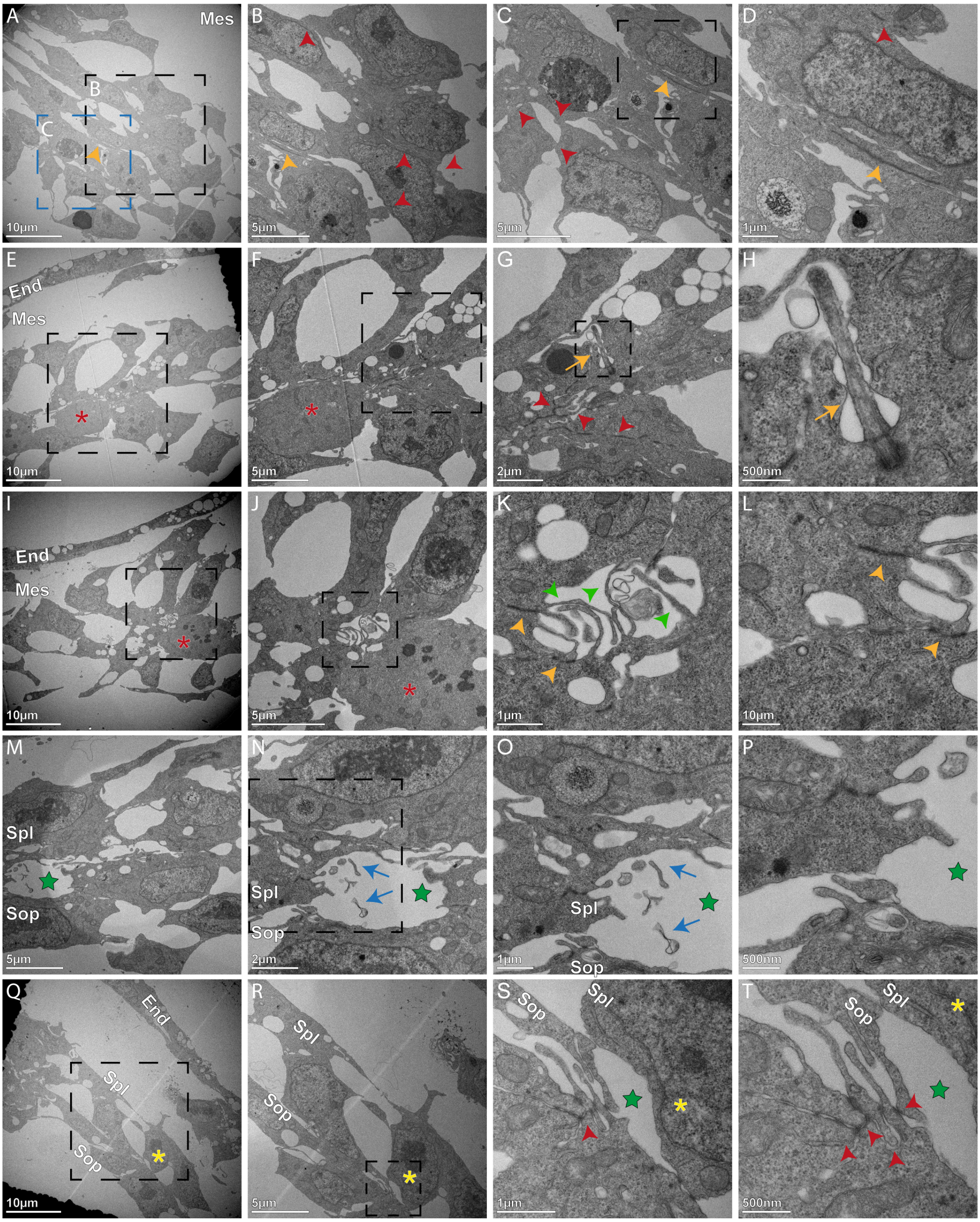


Supplemental Figure S2. Sequential Stages of MET of LPM cells observed by Electron microscopy. In each row the panels are in increasing magnification. At mesenchymal stages (A-D), the mesoderm is multilayered, and the cells exhibit multiple connections with each other (red arrowheads), some via membranal extensions, while others are linearly-arranged (orange arrowheads). At the early rosette stage (E-H), the apical zones of cells are wedge-shaped and closely connected, forming multiple junctional complexes with each other (red arrowheads). There is no coelomic space, and cilia are frequently found in the vicinity of the apical complexes (orange arrow). Asterisk indicates mitotic cell. At the earliest ultrastructural stage in rosette resolution (I-L), a microlumen can be seen containing extensive, thin membrane protrusions (green arrows in K) along with well-developed junctional complexes (orange arrows). Asterisk indicates mitotic cell. M-P show further opening of the lumen (green star), which contains many remnants of membrane protrusions (blue arrows). Q-T show a "hemi-rosette, in which cells on the somatopleuric side are still connected on their apical sides in a rosette-like pattern (red arrowheads) while on the splanchnopleuric side the apical side has broadened into a more mature epithelial phenotype (yellow asterisk). Panels F,G,J,K are the same as Fig. 2E,F,G,H, respectively. End, endoderm; Mes, mesoderm; Sop, somatopleure; Spl, splanchnopleure.

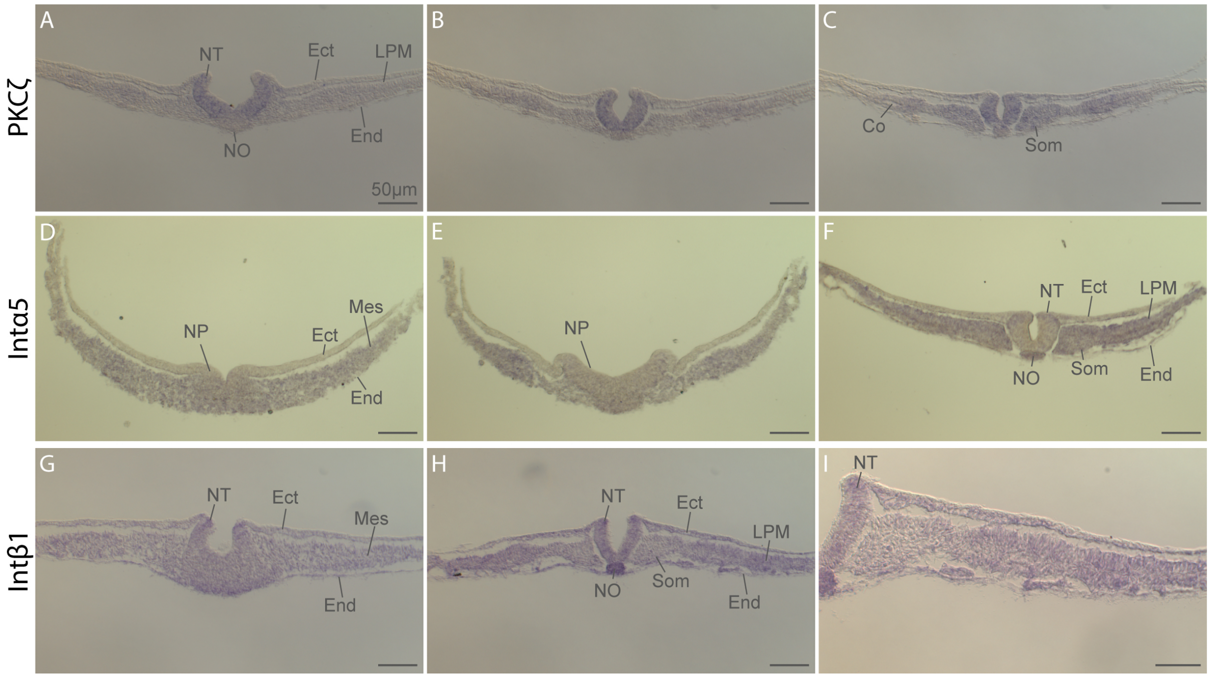


<u>Supplemental Figure S3</u>. <u>PKCζ and Integrin mRNA expression</u>. (A-C) Expression of PKCζ mRNA. Sections of in situ hybridization at successively more advanced stages in the MET process. PKCζ (aPKC) is expressed relatively uniformly throughout the mesoderm, and there is no indication that lateral regions express higher levels of aPKC. (D-I) Expression of Integrin mRNA. Integrinα5 is expressed throughout the mesoderm, but not in the ectoderm. Integrinβ1 is expressed in the mesoderm, ectoderm and endoderm. Ect, ectoderm; End, endoderm; LPM, lateral plate mesoderm; Mes, mesoderm; NO, notochord; NP, neural plate; NT, neural tube; Som, somite.

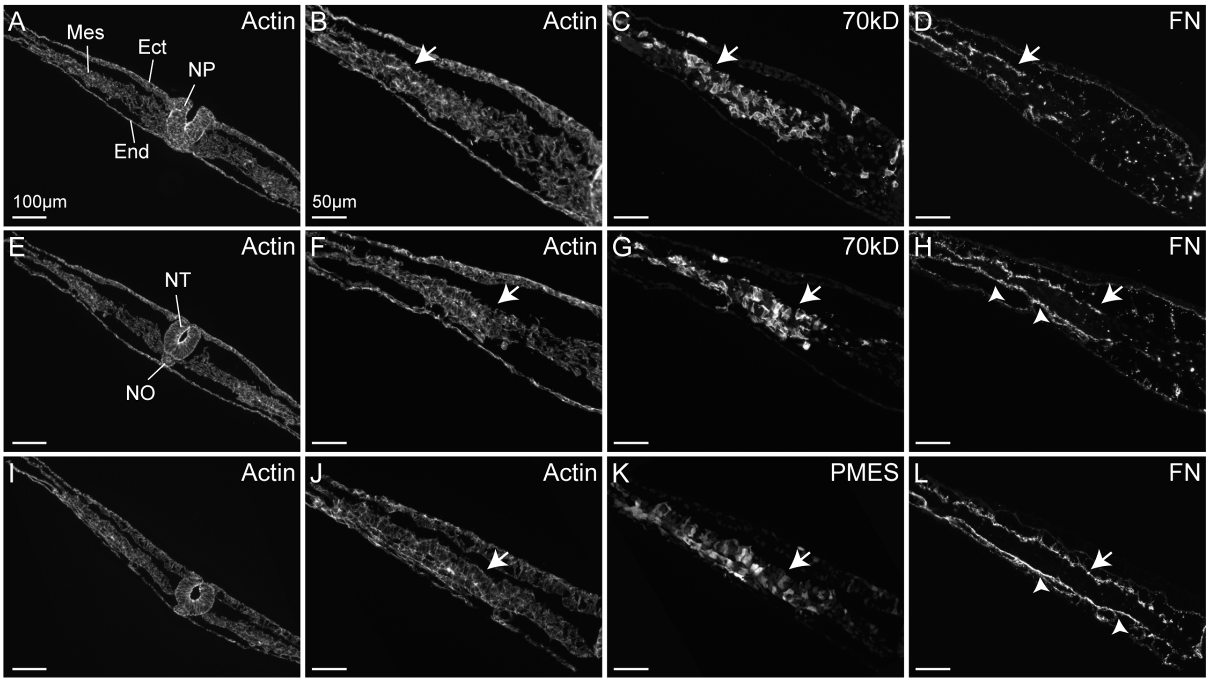


Supplemental Figure S4. MET does not require fibrillar fibronectin. Embryos were electroporated with plasmid coding for a 70kD fragment of FN that interferes with FN polymerization (A-H) or a control plasmid (I-L) and examined at early (A-D) and later (E-L) stages of MET. No difference could be detected in the degree or rate of epithelialization in the presence of the 70kD fragment. Arrows indicate accumulating FN on the basal sides of the forming coelomic epithelium. Arrowheads show the fragmented pattern of FN in 70kD-treated embryos (H), in contrast to the strong and smooth linear pattern in the controls (L), indicating that the 70kD construct prevented normal FN fibril formation. Ect, ectoderm; End, endoderm; mes, mesoderm; NO, notochord; NT, neural tube.

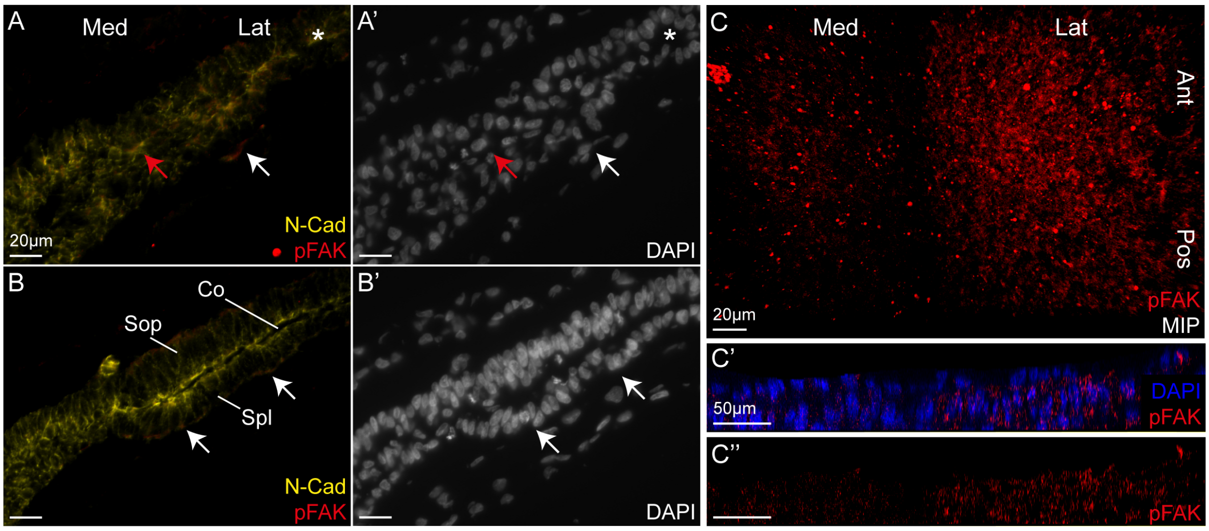


Supplemental Figure S5. Focal Adhesion Kinase (FAK) is activated during MET. Lateral is on the right. (A,B) sections analyzed early (A) and later (B) in the MET process showing presence of phosphorylated FAK (pFAK, arrows) on the prospective apical and basal sides of the coelomic epithelium before epithelialization (red and white arrows in A, respectively) and on the basal side after epithelialization (arrows in B). Asterisk indicates rosette. (C) Whole embryos stained for pFAK and viewed as flat mount in maximal intensity projection (C) or in virtual cross section (C'.C''). Note increased pFAK staining in lateral, more advanced stages of the MET process.

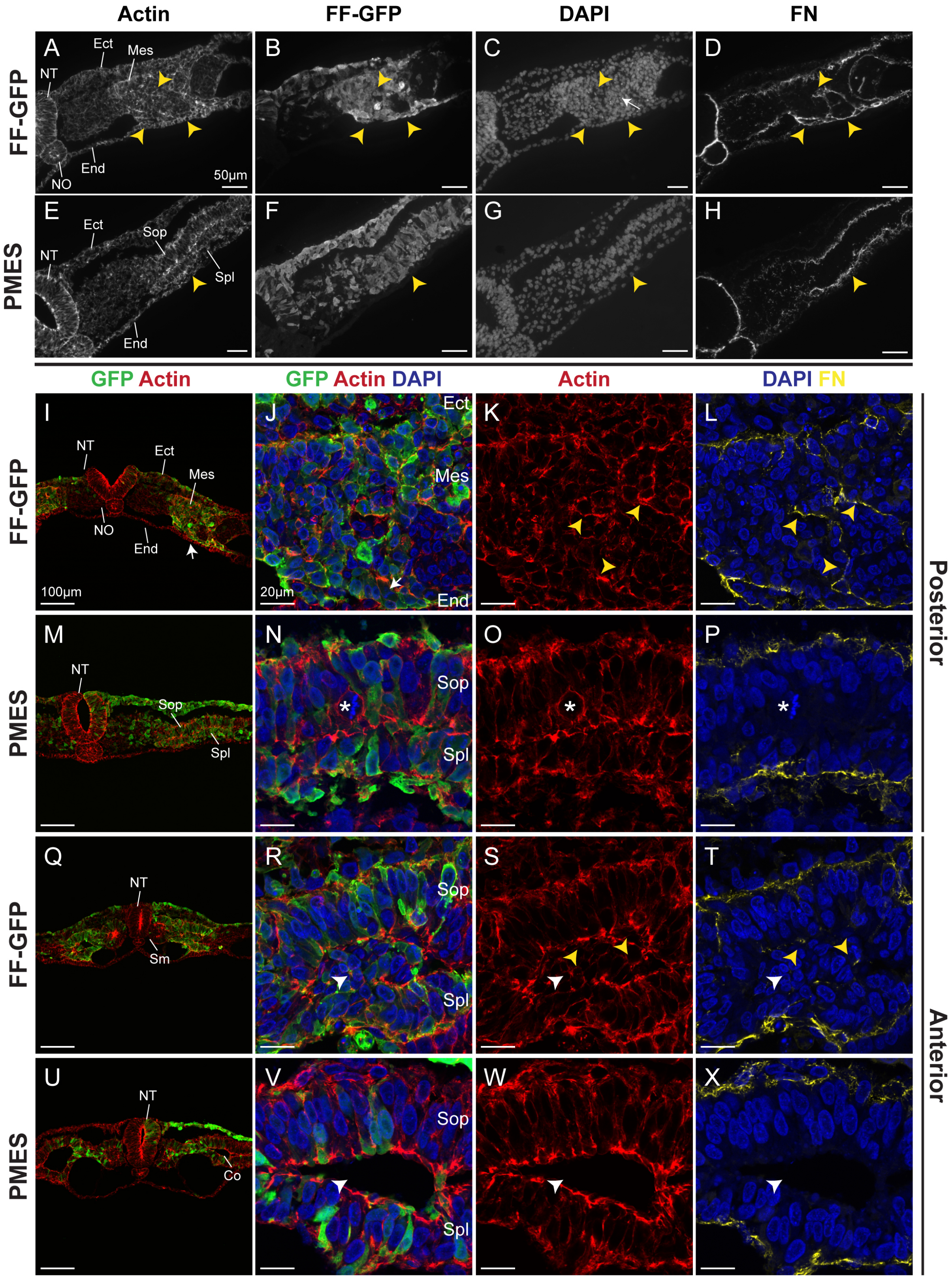


Supplemental Figure S6. Additional data on interference with FAK function, supplement to Figure 5. (A-H) Widefield microscopic images of embryos electroporated with FF-FAK (A-D) or control embryos (E-H). FF-FAK generates large cellular aggregates (yellow arrowheads) surrounded by bands of FN. (I-X) Sections of FF-FAK and control-electroporated embryos at early (I-P) corresponding to (A-H), and later (Q-X) stages of MET. The first panel in each row is a low power view and the remaining panels are higher power confocal images. FF-FAK embryos exhibit cells cast into the coelomic space, clogging the opening (R,S white arrowheads; compare with V,W) and ectopic accumulation of fibronectin (L,T yellow arrowheads; compare with P,X). Ect, ectoderm; End, endoderm; Mes, mesoderm; Sop, somatopleuric mesoderm; Spl, splanchnopleuric mesoderm.

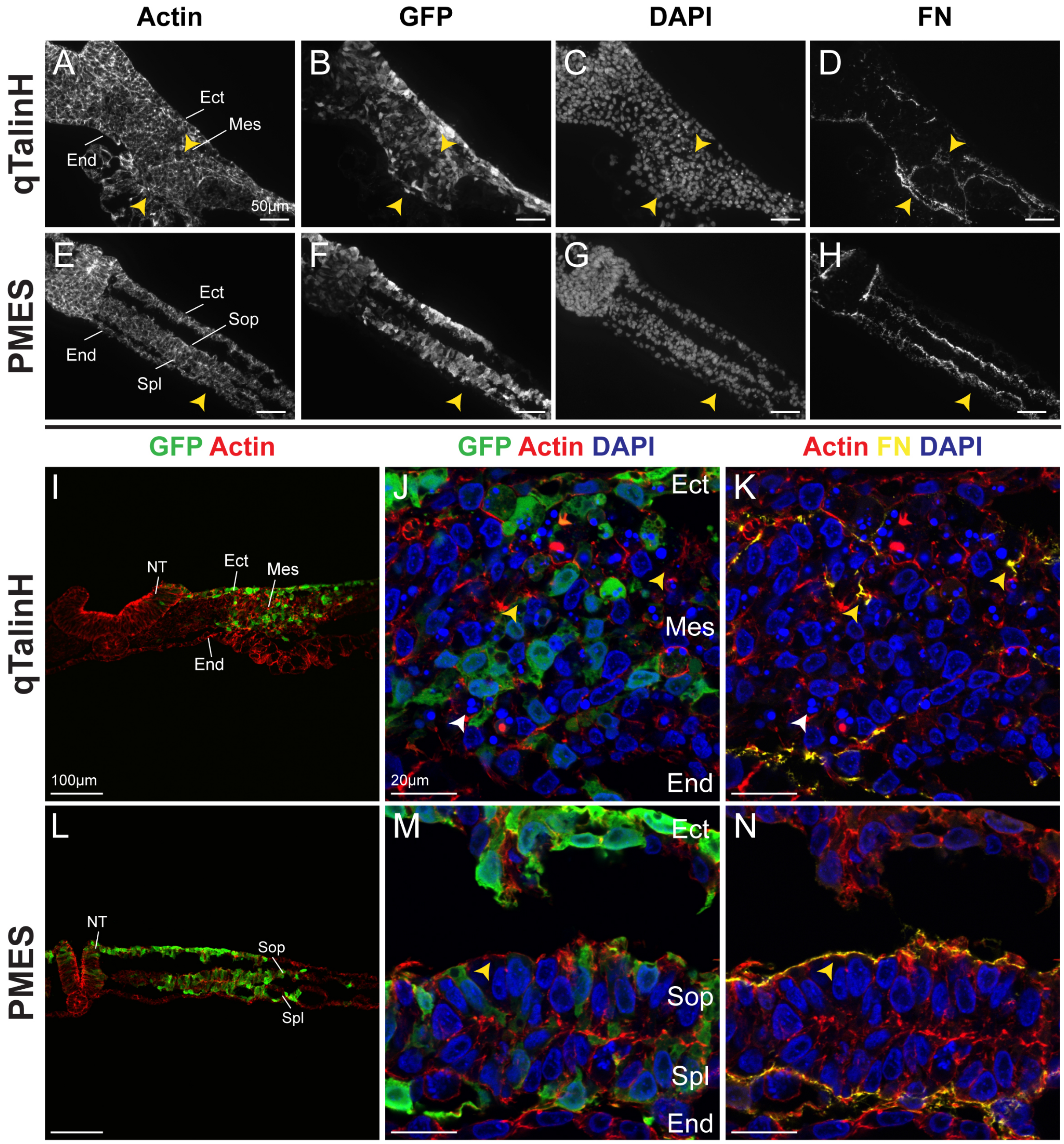


Supplemental Figure S7. Additional data on interference with Talin, supplement to Figure 6. (A-H) Widefield microscopic images of embryos electroporated with qTalinH (A-D) or control embryos (E-H). qTalinH generates large cellular aggregates (yellow arrowheads) surrounded by bands of FN. (I-N) The first panel in each row is a low power view and the remaining panels are higher power confocal images corresponding to A-H. qTalin-electroporated embryos contained disorganized masses of cells which do not undergo epithelialization and exhibit accumulation of fibronectin in ectopic locations (J,K yellow arrowheads) and frequent apoptotic cells (J,K white arrowheads). Ect, ectoderm; End, endoderm; Mes, mesoderm; Sop, somatopleuric mesoderm; Spl, splanchnopleuric mesoderm; NT, neural tube.

Supplemental Videos S1-S3. All videos are 3D segmentation of different regions from the same embryo. Zo-1 is in green and N-Cad is in red. Medial is on the left. Each video starts with whole mount view (XY) of the segmented ROI. Note: Choose download option for high resolution videos.

Supplemental Video S1. 3D Segmentation of the Transition zone. The ROI is located posteriorly. When viewed from the anterior side in cross section (XZ), the rosette is located at the intersection between the unorganized medial side on the left and the two epithelial layers on the right. When viewed again in XY without DAPI, the punctate pattern of Zo-1 and N-Cad on the medial side can be clearly distinguished from the dense staining laterally.
Supplemental Video S1

Supplemental Video S2. Another example of the Transition zone. Here, the rosette can also be seen when viewed in XY. Without DAPI, the punctate pattern of Zo-1 and N-Cad on the medial side is more easily distinguished from the dense staining laterally.
Supplemental Video S2

Supplemental Video S3. The Final stage of Epithelialization and Coelom opening in 3D. ROI is located anteriorly. When viewed from the posterior side in cross section (XZ), two epithelial layers are visible with an open rosette on the medial side. When viewed again in XY without DAPI, the hexagonal pattern of Zo-1 and N-Cad is easily seen.
Supplemental Video S3
